# Linking biotic interactions to species stability

**DOI:** 10.1101/2025.03.25.645234

**Authors:** Ismaël Lajaaiti, Sonia Kéfi, Michel Loreau, Alice Ardichvili, Jean-François Arnoldi

## Abstract

Ecological communities are often composed of many species, each interacting in complex ways. This complexity makes predictions of species responses to disturbances challenging. Here, we analyze dynamical community models and reveal an unexpectedly simple principle: species stability is governed by a single metric—self-regulation loss (SL). SL quantifies the importance of self-regulatory processes in species’ population dynamics. In effect, SL captures a collective outcome of species interactions, organizing how individual species respond to disturbances. When applied to data from protist community experiments, SL accurately forecasts species responses to temperature changes. Our work reveals that, despite the complexity of ecological systems, species stability follows a remarkably simple organizing principle.

## Introduction

Species do not live in isolation: they eat, facilitate and compete with one another (Montoya et al. 2006; Kéfi et al. 2012). Ecological communities consist of vast interaction networks that differ in size, number of links, and structure (Guimarães 2020). These interaction networks strongly influence the population dynamics of the species that form the community (Novak et al. 2016; Bender et al. 1984). For instance, predators feed on herbivores, herbivores deplete plants, and plants compete with one another for limited resources, such that, over ecological time scales, interactions between species mold how biomass flows throughout the community (Patten 1982).

Within communities, species dynamics are not only influenced by direct interactions but also by indirect effects and feedback loops (Abrams et al. 1996; Pichon et al. 2024). For instance, plants depleted by herbivores may benefit indirectly from predators that suppress these herbivores. As interaction networks grow in complexity, indirect effects become increasingly intertwined, eventually reaching a point where the community’s influence on individual species can no longer be understood as a simple sum of direct and indirect interactions (Zelnik et al. 2024). This cohesiveness in community behaviour can produce unpredictable outcomes—rendering species responses to disturbances difficult to decipher (Yodzis 1988; Kawatsu 2024).

Traditionally, studies in functional ecology have emphasized intrinsic species response traits—such as tolerance, avoidance, or regeneration—to understand population stability (Bello et al. 2021; Lavorel et al. 2002). For instance, generation time has been linked to demographic resilience across a wide range of taxa (Gamelon et al. 2014; McDonald et al. 2017; Capdevila et al. 2022). In contrast, community ecology has focused on how interspecific interactions shape the stability of entire communities (May 2019; Kéfi et al. 2019), often through theoretical frameworks such as random matrix theory (May 1972; Allesina et al. 2012; Allesina et al. 2015). However, this focus on the community level often overlooks the variability in species-specific stability (Kéfi et al. 2019). Consequently, the gap between studies that use trait-based approaches to explain stability at the species level and studies that focus on interspecific interactions across entire communities has resulted in a limited understanding of how biotic interactions influence the stability of individual species populations.

A few recent studies have begun to explore species-level stability—examining short-term responses to disturbance (Lajaaiti et al. 2024), non-equilibrium dynamics (Medeiros et al. 2023), or the contribution of individual species to overall community stability (Kunze et al. 2025). Some of these efforts reveal surprisingly simple patterns emerging from complex networks of interactions (Lajaaiti et al. 2024; Kunze et al. 2025). However, many of these efforts either do not explain why such patterns emerge from a mechanistic perspective, or are limited to a single type of disturbance. Here, we build on this work by asking a more general question: can we identify simple, predictive mechanisms by which biotic interactions shape species stability across disturbance types?

Studies focusing on species-level responses have shown that compensatory dynamics can play a key role (Gonzalez et al. 2009; Grman et al. 2010). Compensatory dynamics occur when the decrease in some species abundance is offset by the increase in others. For instance, in plant communities where species compete for limited resources, declines in dominant species biomass following disturbances are often offset by the growth of rare subordinate species benefiting from competition release (Kardol et al. 2010; Mariotte et al. 2013b). These findings highlight the importance of interspecific interactions in stabilizing communities and raise a fundamental question: Is there a general principle organizing species stability across communities, or are these stability patterns so context-dependent that the search for regularities is bound to remain vain?

In fact, although recent work suggests that species stability is linked to their competitive ability (Kunze et al. 2025; Lajaaiti et al. 2024), this relationship is highly contextdependent. The outcome depends on the features of the perturbation. In some cases, compensatory dynamics enhance the stability of subordinate species through competitive release, whereas in others, these dynamics do not occur, and it is the dominant species that are the most stable (Kunze et al. 2025). These findings point to an underlying principle governing species stability within communities. A principle, however, that manifests differently depending on the features of the perturbation (e.g., whether it triggers compensatory dynamics). The precise nature of this principle, its role in shaping stability responses, and whether it exists as a general organizing framework for species stability in complex communities, remains largely unknown.

Starting from a theoretical analysis of dynamical community models, we ask three key questions: (1) Is there a principle that organizes species individual responses to perturbations in complex communities? (2) What stability patterns are expected to emerge from this principle, depending on features of perturbations? And finally, (3) can we validate these predictions using experimental data?

Guided by this line of inquiry, we develop a theoretical framework that reveals the principle organizing species stability across complex communities. Our theory reveals the central role of *self-regulation loss* (SL), a metric which quantifies the relative role of intraspecific regulation processes in species responses. In generalized Lotka-Volterra (gLV) models, this metric coincides with species’ relative yield (RY) (Loreau et al. 2001a), making it empirically accessible.

We show that SL interacts with features of perturbations and determines when compensatory dynamics (and other forms of release in interaction forces) will occur and shape the ordering of species stability within a community. Thus, we could show that the observed context-dependency of stability patterns at the species-level is not at odds with the existence of an underlying, simple, organizing principle.

By integrating theory, simulations and experimental data, our approach not only forecasts the emergence of distinct stability patterns under various disturbance regimes, but also reveals an unexpected simplicity in the organization of species stability. In the following sections, we present our findings, first introducing our theoretical framework and its application to generalized Lotka-Volterra (gLV) models, and ultimately using it to predict species responses to temperature change in a ciliate community experiment (Pennekamp et al. 2018a).

## Methods

### Dynamical model

To study species stability in complex communities, we consider a generalized Lotka-Volterra (gLV) model, where species biomass dynamics follow

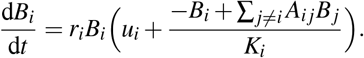

Species in ecological communities interact through interspecific interactions (*A*_*i j*_), which can be either positive (e.g., facilitation) or negative (e.g., competition). For clarity, we focus in the main text on the case where species can persist on their own, thus having growth rates *u*_*i*_ = 1. In this case, *K*_*i*_ corresponds to the carrying capacity of species *i*—that is, its equilibrium biomass when growing alone. We relax this assumption in the Supporting Information (Section S1) and consider obligate species that cannot persist in isolation, in which case *u*_*i*_ *<* 0. Growth rates (*r*_*i*_) set species’ intrinsic timescales, distinguishing faster-growing from slower-growing taxa.

When analyzing species stability near a fixed point, linear interaction terms can be interpreted as local approximations of more complex dynamics, including nonlinear and higher-order interactions (for generalization to nonlinear models see SI Section S1).

In the following, we derive expressions for species responses to changes in environmental conditions (press disturbances) and to biomass removal events (pulse disturbances). We show that these responses can be succinctly expressed as a function of species self-regulation loss SL, which for Lotka-Volterra models and when species can grow on their own, coincides with species’ relative yields *B*_*i*_*/K*_*i*_ (Box 1 and Fig. 1B).

**Figure 1:**
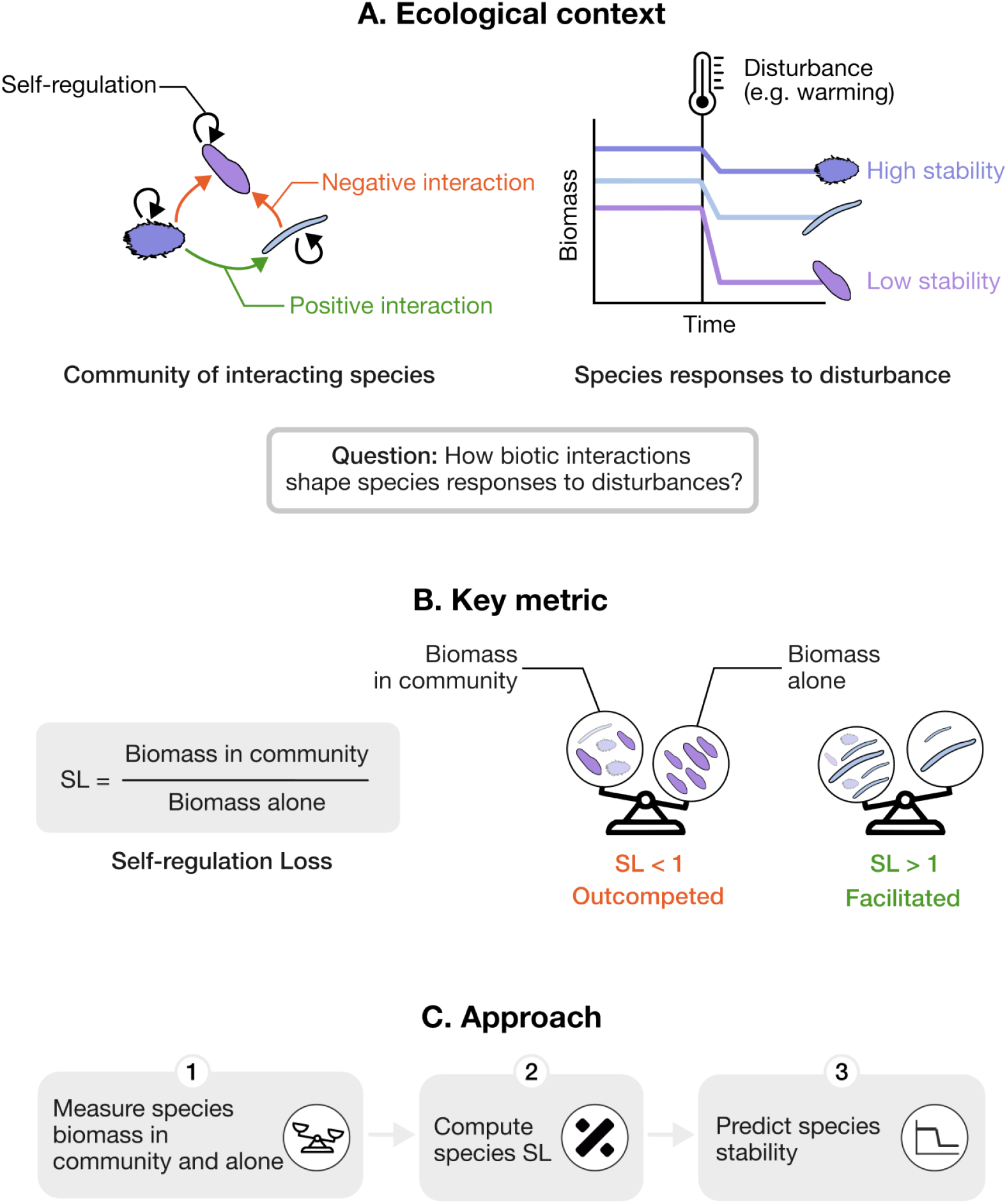
Linking biotic interactions to species stability through self-regulation loss (SL). A) Species within ecological communities interact in both positive and negative ways. When a disturbance occurs (such as warming), species respond differently—some are stable, others less so. This variation is partly explained by traits (such as thermal niche), but here we ask: how do biotic interactions shape these responses? B) We show that a single quantity—self-regulation loss—captures how biotic interactions influence species stability. C) SL can be estimated from simple biomass measurements (alone and in community), providing a tractable way to predict how species respond to disturbance based on how they interact with the rest of the community.

### Derivation of species response to press

At equilibrium, species biomass satisfies

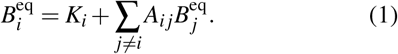

In the absence of interactions, species biomass in the community equals their carrying capacity. However, as interaction strength increases, species biomass becomes increasingly shaped by the biotic environment (Zelnik et al. 2024). By rearranging Eq. 1, we isolate the carrying capacity:

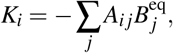

where *A*_*ii*_ = −1. In vector form, this equation reads

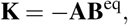

where **K** is the vector of species carrying capacities, **B**^eq^ is the equilibrium biomass vector, and **A** is the interaction matrix.

Defining the sensitivity matrix as **V** = −**A**^−1^, we can express equilibrium biomass as

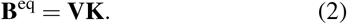

Now, we introduce a press disturbance, modeled as a change in the carrying capacity given by the vector Δ**K**. The resulting change in equilibrium biomass follows

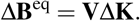

To quantify how biotic interactions influence species stability, we define species sensitivity to press disturbances as the ratio of its relative biomass change in the community to its relative biomass change in isolation. This definition ensures a fair comparison across species with different biomasses or carrying capacities. Biomass changes in isolation and in the community are given by

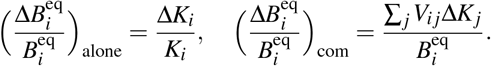

Taking their ratio, species sensitivity is given by

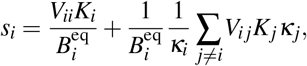

where 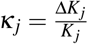 denotes the press intensity.

In disordered communities where interactions are weak enough to permit stable coexistence, the diagonal elements of the sensitivity matrix are approximately equal across species and close to one, *V*_*ii*_ ≃ 1 (but see Supporting Information, Section S4). To further simplify, we adopt a mean-field approximation, assuming that press intensities *κ*_*j*_ are weakly correlated with the net interaction terms *V*_*i j*_*K*_*j*_ leading to

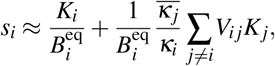

where 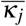 is the average press intensity across species except *i*. Using Eq. 2 for species *i*, and introducing species self-regulation loss 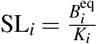 we obtain the following expression for species sensitivity:

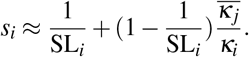

Finally, taking the expected sensitivity across many press experiments yields

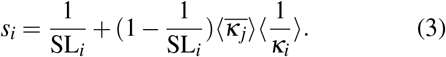

For antagonistic release to prevail, the disturbance must impact other species sufficiently; specifically, when 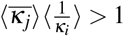. This condition is typically met when disturbances affect all species similarly, as in this setting arithmetic mean exceeds harmonic mean (that is, 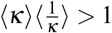.

### Derivation of species responses to pulse

We now focus on species responses to pulse disturbances, which we model as an instantaneous perturbation in biomass, 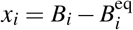. Assuming small deviations from equilibrium, we linearize the system around 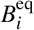, leading to the species recovery equation, normalized by its generation time 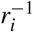 as suggested by Mentges et al. 2024:

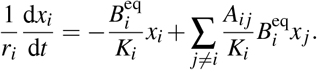

Introducing species self-regulation loss (SL) and the relative biomass deviation 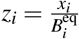, we obtain

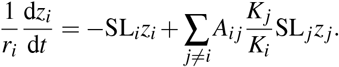

To focus on interactions shaping species coexistence, we introduce core interactions 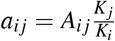. These rescaled interactions are the relevant terms that determine species coexistence (Barbier et al. 2018).

Following a mean-field approximation as before, we assume weak correlations between *a*_*i j*_SL _*j*_ and *z* _*j*_, a reasonable assumption in species-rich communities, when initial perturbations *z* _*j*_(*t* = 0) are independent of species interactions. This simplifies the equation to

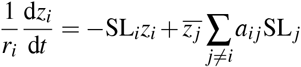

where 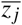 denotes the mean species deviation. Using Eq. 1 (divided by *K*_*i*_), we simplify the sum to

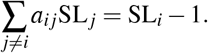

Substituting this into the previous equation defines an effective model for the dynamics governing species recovery

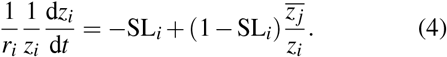

To solve this equation, we first derive the trajectory of the average species deviation, 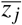. We assume that 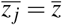, the mean deviation over all species, including *i*. It can then be shown that the trajectory of 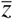 follows a simple exponential decay:

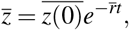

where 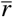 is the average species growth rate. This allows us to express the recovery trajectory of species *i*:

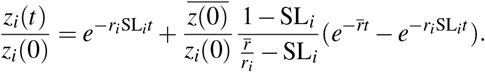

Taking the expected value over multiple disturbances (Arnoldi et al. 2018), denoted by ⟨·⟩, and assuming *z*_*i*_(0) follows a common distribution with arithmetic and harmonic means ⟨*z*_0_⟩ and 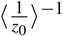, respectively, we obtain

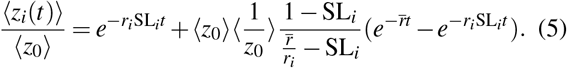

From this, we derive the expected species average recovery rate between 0 and *t*:

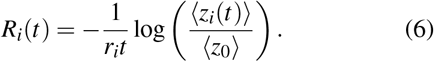

where *z*_*i*_(*t*) is given by Eq. 5. This full expression underpins our analytical predictions, but further approximations allow for better interpretation. In the short term, the recovery rate simplifies to

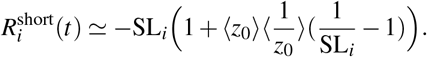

This demonstrates that species SL dictates two fundamental aspects of species recovery: their characteristic recovery timescale and the strength of antagonistic release. Specifically, we observe that a species’ recovery rate is proportional to its SL. Additionally, the term capturing antagonistic release is proportional to 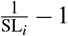, meaning it is large when SL_*i*_ ≪ 1 and vanishes when SL_*i*_ = 1. However, in nonlinear models where SL is no longer equivalent to RY, these two recovery features are not solely controlled by SL (Fig. S1). While the characteristic recovery rate remains governed by SL, antagonistic release is instead determined by RY.

In the long term, antagonistic release vanishes, leaving only species characteristic recovery rate to govern their recovery:

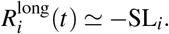

### Simulations

We simulated communities of *S* = 30 species. Following previous studies (Barbier et al. 2018; Bunin 2017), we draw core interspecific in Gaussian distributions 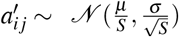, with *µ* = −1 and *σ* = 0.3. Furthermore, *a*_*ii*_ = −1 in our parametrization as self-regulation is already captured by species carrying capacity *K*_*i*_, that are drawn in a Gaussian distribution *K*_*i*_ ∼ 𝒩 (1, 0.3). To center on the effect of interspecific interactions on species stability, we took *r*_*i*_ = 1 for all species. This choice has no consequence for press disturbances, as species’ changes in biomass—and thus their sensitivity—do not depend on their growth rates (Eqs.2, 3). For pulse disturbances, we explore in the Supporting Information (Section S9) how variation in species’ growth rates affects the accuracy of our predictions. Press intensities were drawn in *κ* ∼ LogNormal(log 0.1, 0.5), and pulse disturbances in *z*_0_ ∼ LogNormal(log 0.01, 0.65) for all species. Simulations were run in the Julia programming language (Bezanson et al. 2012) using the DifferentialEquations package (Rackauckas et al. 2017).

### Data

We analyzed data from a previously published experiment (Pennekamp et al. 2018a), which factorially manipulated temperature (15, 17, 19, 21, 23, and 25 °C) and species richness (1–6 species) in assemblages of bacterivorous ciliates: *Colpidium striatum* (Colp), *Dexiostoma campylum* (Dexio), *Loxocephalus sp*. (Loxo), *Paramecium caudatum* (Para), *Spirostomum teres* (Spiro), and *Tetrahymena thermophila* (Tetra). Species biomasses were measured over a period of 40 days.

#### Carrying capacity

We estimated species carrying capacity (*K*_*i*_) from their biomass when grown alone (monoculture).

#### Sensitivity

To quantify species sensitivity in the experiment, we expressed it as:

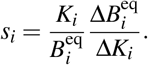

Here, *K*_*i*_ represents the mean carrying capacity observed across all temperatures, while 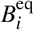 is the equilibrium biomass of species *i* across temperatures for each community composition. To minimize transient effects, we restricted analyses excluded the first seven time points from the time series. Moreover, we considered communities with at least three species.

Temperature changes affected species equilibrium biomass both in isolation (carrying capacity) and within communities. We estimated the term 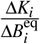 as the slope of a linear model relating species biomass in communities 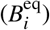 to their carrying capacity (*K*_*i*_) (see Fig. 4B,C). This analysis was performed for each species and community combination, making species sensitivity specific to both the species and its community context.

Our analytical prediction of species sensitivity (Eq. 3) depends on species’ SL, but also on the arithmetic and harmonic means of press intensities on species (*κ*_*i*_). We estimated the press intensity as

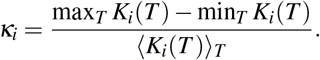

which gives us the relative change in species’ carrying capacity induced from temperature change, across the gradient considered.

#### Interspecific interactions

We inferred pairwise interactions from species biomass when grown in duocultures for each temperature treatment. From Eq. 1 we have that

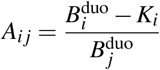

which give us species interactions based on their biomass in mono and duocultures made of species *i* and *j*. From Eq. 1, we expect interactions to shape species SL such as

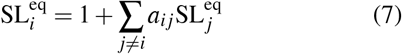

where we have switched to the core *direct* interactions 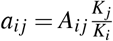.

From this, we computed the incoming direct interactions a species receives, given by ∑ _*j*≠ *i*_ *a*_*i j*_. This sum indicates whether a species is predominantly influenced by negative or positive interactions. However, we do not expect it to perfectly correlate with species SL. As shown in Eq. 7, this sum is effectively weighted by the SL of the interacting species. These weights capture how the focal species influences the biomass of those it interacts with, leading to a measure of incoming net effects. Formally, this relationship can be derived by inverting Eq. 7. We thus expect incoming net interactions to show a stronger correspondence with the observed SL values.

## Results

We asked how biotic interactions shape species stability in the face of disturbances (Fig. 1A). Through an analytical approach, we found that a single quantity—selfregulation loss (SL)—captures the effect of interactions on how species respond to perturbations (Fig. 1B). SL is captured by a simple biomass ratio: a species’ biomass in community relative to when it grows alone. This one number reflects how the biotic environment shapes the species—whether it’s helped, hindered, or unaffected by others (see Box 1 for details). In the following, we test in simulated communities how SL predicts species’ responses to long-lasting changes in environmental conditions such as warming (press), and shock events such as biomass removal (pulse). We moreover validate our predictions on experimental data from protist communities exposed to warming (Fig. 1C).

### Predicting response to press

We modelled press disturbances by altering species carrying capacities. Following a press disturbance, species biomass shifts, and the magnitude of this change reflects the species’ sensitivity to the disturbance. We defined sensitivity as the ratio of the relative change in biomass observed in a community to the change if the species was alone (Methods). In this way, our measure captures how interspecific interactions influence the species’ stability and we further ensure a fair comparison among species.

#### Single-species press

We first considered the simplest scenario, where only a single species, *i*, is affected by a press perturbation, and we focus on the sensitivity of that species only. In this setting, the contribution of other species is null as they are not directly disturbed. As a result, species sensitivity equates the inverse of its SL (Eq. 3 with *κ*_*j*_ = 0).

To test this prediction, we assembled a large generalized Lotka-Volterra (gLV) community and measured each species’ SL before applying any disturbance. We then proceeded by applying targeted disturbances on each species successively and measured their resulting sensitivity. Our prediction closely matched observed species sensitivity (Fig. 2B).

**Figure 2:**
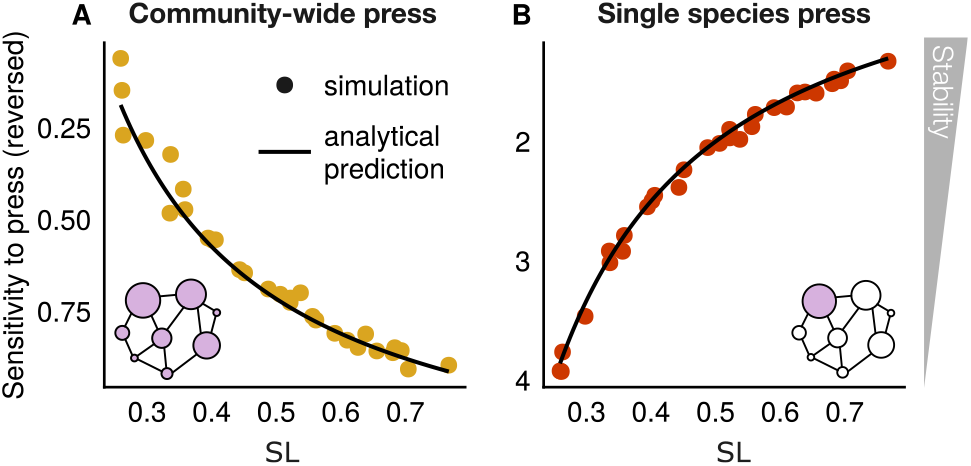
Species’ self-regulation ratio (SL) organizes their stability to press disturbances. (A) Sensitivity of species to communitywide press perturbations, where the press is applied simultaneously to all species (Methods). Sensitivities are averaged over 1,000 disturbance events. Analytical prediction is given by Eq. 3. Sensitivity of species to single-species press perturbations, where the press is applied to one species at a time (Methods). Sensitivities are averaged over 50 disturbances per species. Analytical prediction is given by Eq. 3 taking ⟨*κ*_*j*_ = 0⟩. Note that the sensitivity-axis is reversed so that upper points indicate greater stability.

The inverse relationship between species sensitivity and SL arises because a species’ biomass change under press disturbance closely follows the change in its carrying capacity. Importantly, interactions don’t directly buffer or amplify the press disturbance itself. Instead, they shape a species’ baseline biomass, which then determines how strongly the species responds.

If a species has an SL of one, it means that interactions have little effects on its biomass—so its sensitivity is also close to one. If interactions suppress a species (SL *<* 1), it start from a lower biomass and experience a proportionally larger loss, leading to a sensitivity above one. If interactions boost a species (SL *>* 1), it start from a higher biomass and show a smaller relative loss, leading to a sensitivity below one.

When we examined the response of species that are not directly targeted by the press disturbance, the pattern reverses (see SI Section S7). Species with higher SL are less stable than those with lower SL. This is because their response is dominated by interaction release. Facilitated species with high SL are more vulnerable to disturbances affecting their interacting partners, since their biomass relies heavily on those positive interactions. In contrast, species burdened by strong antagonistic interactions benefit from the disturbance of others, as they experience a release from antagonistic pressure.

When a press perturbation targets a single species, its response is shaped by self-regulation: the higher its SL, the more stable (i.e., less sensitive) it is. In contrast, other species respond through interaction release: the lower their SL, the more stabilizing the release effect is, and the less sensitive they are. In community-wide press scenarios, species responses are shaped by both mechanisms—selfregulation and interaction release—leading to a more nuanced stability pattern.

#### Community-wide press

While targeted disturbances affect a single species, environmental changes often impact entire communities. We next tested our predictions in the scenario of community-wide press, where all species are affected by the disturbance. Under this scenario, we found that species sensitivity is also shaped by the response of others, which adds a new term to the expression of species sensitivity (Eq. 3). Remarkably, the net impact of the community on species sensitivity can be expressed as a simple function of SL.

##### BOX 1 Key concept: self-regulation loss (SL)

**Motivation** When a community of interacting species experiences a disturbance, individual species often respond in different ways (Fig. 1A). Beyond traits, a key source of this variability is how species dynamics are shaped by interspecific interactions. Yet, interactions are numerous, rendering their effects hard to disentangle. We approach this task by deriving analytical expressions of species responses to disturbances (Methods). These derivations reveal a key term that appears to govern species responses: self-regulation loss (SL).

**Definition** SL measures the biomass loss due to self-regulation relative to intrinsic growth. By adjusting for intrinsic growth, it becomes a unitless measure, making it comparable across species with different generation times. In the gLV model, SL writes as the ratio of a species’ biomass in the community to its biomass in isolation. This aligns with the classic concept of relative yield (RY) (Loreau et al. 2001b), offering an intuitive interpretation of SL. However, this correspondence does not generally hold in more complex models (Fig. S1). At equilibrium, SL writes

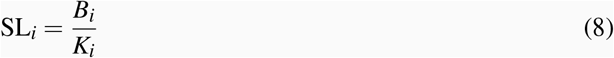

**Interpretation** SL reflects how a species’ biomass is influenced by its interactions with other species. Species experiencing strong competition have low SL, while those benefiting from facilitation show high SL. Notably, species benefiting from strong facilitation can surpass their carrying capacity, leading to SL values greater than one. In this view, although SL captures the degree of self-regulation a species experiences, it is also shaped by interspecific interactions. For instance, the same species may show higher self-regulation in a community where it is facilitated and maintains high density, compared to one where it is outcompeted and persists at low density.

**In practice** Self-regulation loss (SL) is ideally evaluated when species are at equilibrium, since deviations from equilibrium biomass can lead to over- or underestimation. This assumption can limit the direct application of SL to empirical data, where equilibrium is rarely observed. However, we show that SL is related to metrics used to predict species sensitivity under non-equilibrium dynamics (see SI Section S8). While calculating SL requires knowledge of species’ density dependence (e.g., biomass when grown in isolation), it does not rely on detailed measurements of all pairwise interaction strengths. This is a key advantage, as such interactions are often difficult to estimate reliably and are highly sensitive to observational error (Terry 2025). By sidestepping the need for a fully specified interaction network, SL offers a more robust and empirically tractable approach to understanding species-level stability.

##### BOX 2 Self-regulation Loss in practice: a two-species illustration

To build intuition for how self-regulation loss (SL) predicts species-level stability, we consider a simple community of two species. To ease interpretation, we assume that species 1 is unaffected by species 2, while species 2 is influenced by species 1. The dynamics are given by

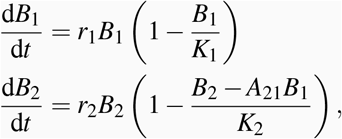

where *K*_*i*_ is the carrying capacity and *a*_21_ captures the effect of species 1 on species 2.

We apply a press disturbance, modeled as a proportional reduction in carrying capacity: Δ*K*_*i*_ = −*κ*_*i*_*K*_*i*_, where *κ*_*i*_ *>* 0 indicates disturbance intensity. The change in equilibrium biomass is

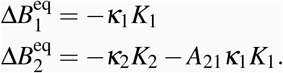

To compare responses across species, we compute their sensitivity by normalizing the biomass change by equilibrium biomass and press intensity:

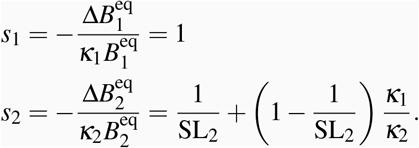

Here, sensitivity *s*_1_ = 1 reflects that species 1 is unaffected by species 2 and behaves as if in isolation. This provides a useful baseline: when interactions are absent, the species response in community equals its response when grown alone.

By contrast, species 2 is influenced by interactions. When *κ*_1_ = 0, its sensitivity simplifies to 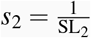. This means the more species 2 benefits from interactions (higher SL), the more buffered it is against disturbance. For instance, a facilitative effect (*A*_21_ *<* 0) stabilizes species 2, reducing *s*_2_.

However, when both species are disturbed facilitative effect can become destabilizing, because of the release of facilitative interaction following the disturbance. Specifically, when species 1 is more affected (*κ*_1_ *> κ*_2_), interaction release dominates. The sensitivity of species 2 *increases* with its SL reflecting the destabilizing effect of facilitative release (if *A*_21_ *>* 0), or the stabilizing effect of antagonistic release (if *A*_21_ *<* 0).

A species’ response to disturbance is shaped not only by its self-regulation, but also by interaction release (captured by the second term in Eq. 3). When all species are similarly affected by a disturbance, interaction release dominates. In this case, species with SL*<* 1—those suppressed by antagonistic interactions—exhibit greater stability due to the buffering effect of antagonistic release, leading to sensitivities below one. In contrast, species with SL*>* 1 lose facilitative interactions and become less stable.

### Predicting response to pulse

Having established that the SL structures species stability under press disturbances, we next sought to find whether these findings extend to pulse disturbances—sudden, temporary shifts in species biomass. We quantified species stability through their recovery rate, which describes how quickly species return to equilibrium after the disturbance (Methods).

We studied the recovery of species relative deviation around their equilibrium. To do so, we linearized the dynamics (assuming small deviations), and we used a meanfield approximation similar as the one performed for press disturbance (Methods). This simplifications reduces the recovery dynamics to two key components (Methods; Eq. 4).

The first term reflects the species’ ability to recover as if it was the only species perturbed, that is, its characteristic recovery rate. The second term captures the contribution of the community to the species stability. High-SL species benefit from strong self-regulation, while low-SL species gain stability from interspecific interactions—provided that the disturbance affects the species they interact with. These two mechanisms echo those we previously identified for press disturbances.

Using an approach similar to (Arnoldi et al. 2018), we solved Eq. 4 to derive the expected recovery trajectories over a large set of pulse disturbances. This analysis revealed broad, recurring patterns across disturbances, notably uncovering two distinct phases of recovery (Fig. 3A). Immediately following a pulse disturbance, species with low SL recover rapidly due to antagonistic release triggered by the decline of their suppressors. This rapid recovery, however, is followed by an overshoot of equilibrium biomass—as these species reach equilibrium while still benefiting from antagonistic release. As the community approaches equilibrium and interspecific effects wane, intrinsic resilience takes over, ultimately favoring species with high SL.

**Figure 3:**
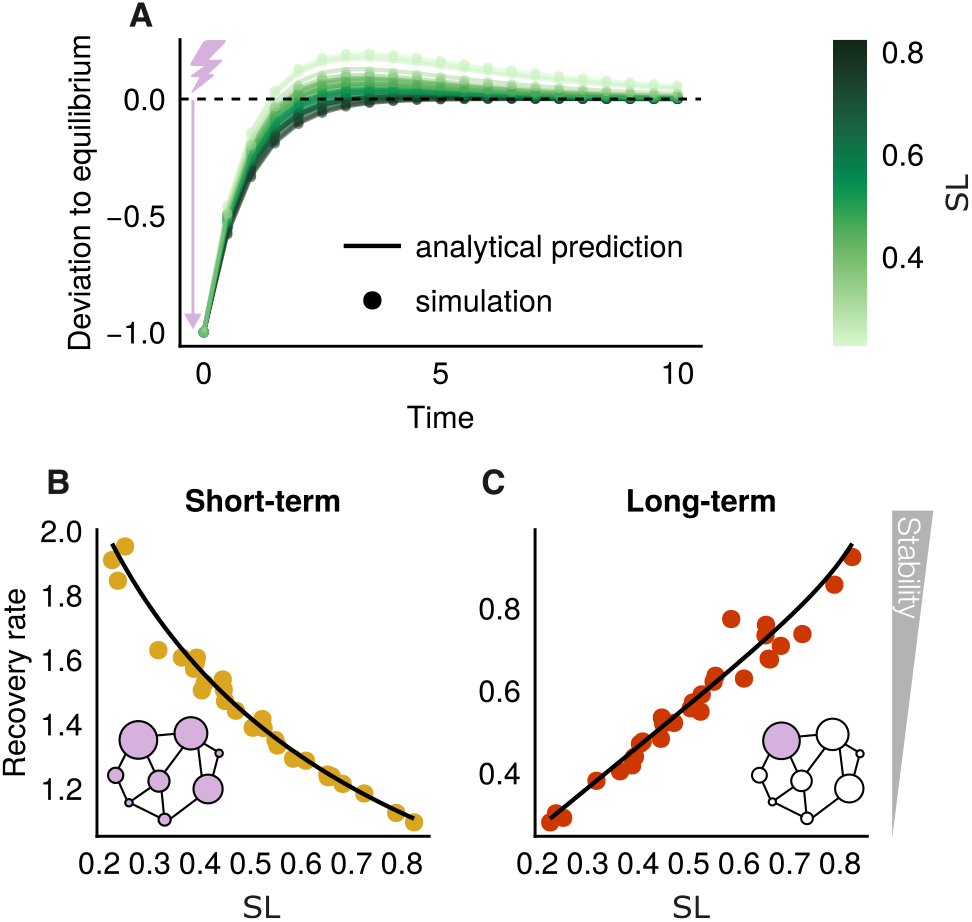
Species’ SL organizes their recovery to pulse disturbances. (A) Species recovery trajectories over time. Trajectories are averaged across 5,000 random pulse perturbations (Methods). Deviation to equilibrium is defined as 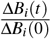 with 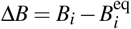, normalized so all species start at a value of −1. Over time, species return to equilibrium, with the deviation approaching 0. Analytical prediction is given by Eq. 5 (Methods). (B) Short-term return rate vs. SL. The return rate is averaged over the interval *t* = 0 to *t* = 1. The network indicates that in the short-term all species are far from their equilibrium, making the link with the stability pattern observed for community-wide press (Fig. 2A). (C) Long-term return rate vs. SL. The return rate is averaged over the interval *t* = 0 to *t* = 10. Networks do not reflect the initial extent of the disturbance, as shown in Fig.2; in both cases, the pulse affects all species. However, they illustrate that in the short term, all species deviate significantly from equilibrium, whereas in the long term, only a few have not recovered, linking this pattern to the stability responses observed in single-species press disturbances (Fig.2B).. Averages are computed over 1,000 random pulses. Analytical prediction for the recovery rates is given by Eq. 6 (Methods).

**Figure 4:**
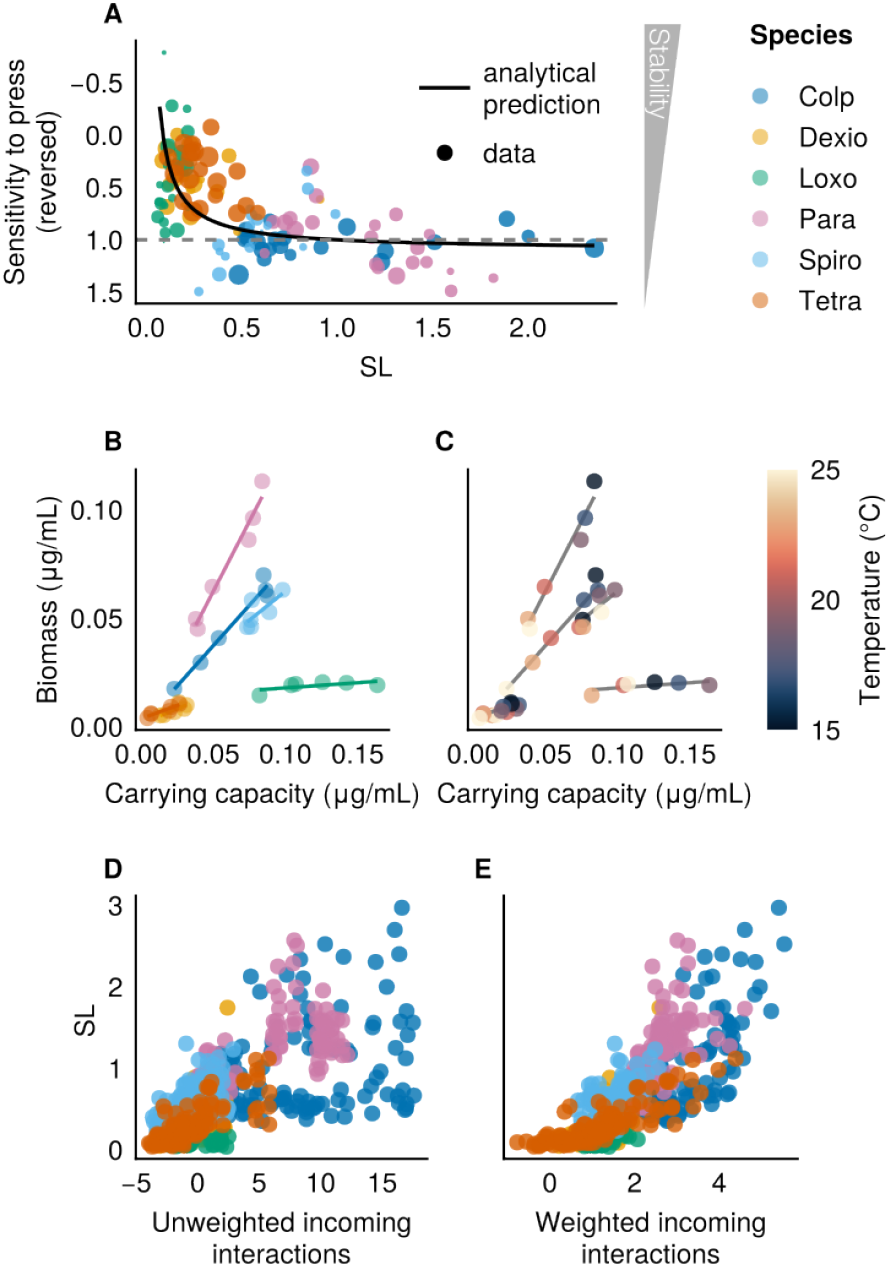
Predicting the response of ciliate communities to temperature change. (A) Species sensitivity to temperature change against their SL, with the analytical prediction based on Eq. 3 (Methods). The grey, dashed line gives the baseline *s* = 1, which is expected if species do not interact. That is, if their response to warming is fully determined by their response in monoculture, and so, their intrinsic features. Point sizes decrease with the size of the confidence interval. (B) Species average biomass in community against their carrying capacity colored by species. (C) Same plot but colored by temperature to show the impact of temperature on species biomass and carrying capacity. As temperature increases, both parameters tend to decrease. (D) Species SL against the incoming direct interactions inferred from duocultures (see Methods). (E) Same but the incoming *net* interactions that account for the effect of the focal species on others.

This behavior is captured quantitatively by examining species’ short- and long-term average recovery rates (Methods). In the short term, species with low SL recover faster because of antagonistic release (Fig. 3B). Over longer time scales, however, the advantage of antagonistic release diminishes, and species with high SL—benefiting from strong self-regulation—recover more rapidly (Fig. 3C).

### Empirical validation

We next tested the applicability of our framework on experimental data. We reanalyzed time series data from (Pennekamp et al. 2018a), which monitored ciliate communities over 57 days. These communities were assembled with varying species compositions (ranging from one to six species) and were exposed to a temperature gradient from 15 to 25°C.

In our analysis, we defined the press disturbance as the temperature change that alters species biomass both in isolation and within the community (Fig. 4B,C). We observed a linear relationship between species biomass in the community (averaged on all community combinations) and their carrying capacity, which validates a key assumption of our framework.

We further estimated the sensitivity of each species in each community composition (of three species or more) by fitting a linear model on their biomass in this community and their carrying capacity, across the range of temperatures studied (Methods). Because species were also grown in monoculture, we were able to infer species carrying capacity (Methods). From this, we were able to quantify species SL. As a result, we obtained how species sensitivity to temperature change depends on their SL in each community composition. We tested our analytical predictions (Eq. 3) against these data. Remarkably, our prediction closely captured the observed species sensitivities to temperature change (Fig. 4A).

A species’ SL is not fixed—it depends on the community context. Changes in species composition and interaction strengths can shift a species’ SL, underscoring how biotic environment shapes stability. To empirically establish the direct connection between a species SL and its biotic environment, we estimated pairwise interactions from duoculture biomass data (see Methods). These estimates allowed us to quantify the total incoming interactions a species receives (Fig. 4D,E). The unweighted sum captures the overall balance of positive and negative effects acting on a species. The weighted version goes further, adjusting for how the focal species influences the biomass of its interaction partners (see Methods).

We found a strong positive relationship between incoming interactions and observed SL values. This relationship became clearer when we accounted not only for how other species affect the focal species, but also for how the focal species influences others. These results confirm that SL captures key aspects of a species’ biotic environment.

Importantly, the variability in species sensitivity is well-captured by its SL. This demonstrates that species sensitivity is not solely the result of its intrinsic properties (e.g. a life-history trait) but is also shaped by the community it lives in.

If species sensitivity were determined solely by their intrinsic response to temperature change, without being influenced by interspecific interactions, sensitivity would equal one (grey line Fig. 4A). In this case, a species’ response to temperature changes would be the same in both the community and in isolation, resulting in a sensitivity of one. Therefore, the observed decrease in sensitivity for species with low SL indicates that interactions do affect species stability. Specifically, species with low SL are stabilized through antagonistic release.

Notably, we observed ciliates with SL values largely exceeding one, indicating the presence of facilitative interactions (e.g., *Dexiostoma campylum*, Dexio, in Fig. 4A). Our prediction works well across the entire range of observed SL, demonstrating that the mechanisms it captures are not limited to competitive communities but also apply to communities with more positive interactions (see also SI Section S6). Our framework explains the higher sensitivity (lower stability) of species with high SL due to the destabilizing effects of facilitative interaction release, while low-SL species gain stability from antagonistic release. Together, these findings highlight key mechanisms driving species stability within real ecological communities.

The stabilizing effect of antagonistic release (when SL *<* 1) and the destabilizing effect of facilitative release (when SL *>* 1) are not symmetrical. The stabilizing influence of antagonistic release is markedly stronger than the destabilizing effect of facilitation. This asymmetry is reflected in the way species sensitivity saturates around 1 as SL increases (Fig. 4).

## Discussion

Understanding how interspecific interactions shape species stability is a threefold challenge. First, interactions are numerous, and their effects on species dynamics are often difficult to decipher (Zelnik et al. 2024; Kawatsu 2024; Yodzis 1988). Second, it is practically challenging to measure all pairwise interactions within a community (Carrara et al. 2015; Lubiana Botelho et al. 2025), assuming that pairwise interactions still make sense when considering more than two species (Battiston et al. 2021; Abrams 1983). Third, species stability is highly context-dependent and may not exhibit consistent patterns across different communities and perturbation scenarios. Here, we show that the challenge of understanding species stability in complex communities can be addressed using a single species-level metric: self-regulation loss (SL). This metric captures how biotic interactions shape species stability.

While SL quantifies self-regulation, it is not only determined by a species’ intrinsic properties. It also reflects the broader community context. This captures a simple but important intuition: a species suppressed to low density by negative interactions will exhibit lower self-regulation than the same species thriving in a more favorable community.

A key strength of our approach lies in the simplicity of SL—a biomass ratio that captures how a species is shaped by its biotic environment. Crucially, SL does not require reconstructing the full interaction network, which is often infeasible in practice. As noted by Terry (2025), avoiding reliance on exact network structure helps reduce sensitivity to observation error. This works because a species’ equilibrium biomass already reflects its biotic context. The assumption of equilibrium is thus not essential per se for the framework to hold—what matters is that a species biomass reflects the effects of interactions.

When disturbances reduce species biomass, they also weaken the effect of biotic interactions. This interaction release can lessen antagonistic pressures like competition or predation, helping to buffer species from further biomass loss. Such dynamics are known as compensatory dynamics (Gonzalez et al. 2009). In climate change experiments, compensatory dynamics have been shown to stabilize plant communities during drought, as rare species increase in biomass to offset the decline of dominant ones (Kardol et al. 2010; Mariotte et al. 2013a).

We identify a simple condition under which interaction release—and thus compensatory dynamics—becomes the main driver of species stability. This occurs when the species interacting with a focal species are disturbed as much or more than the focal species itself. This for example occurs in communities where all species experience similar levels of disturbance. This results is inline with recent results on species contribution to community stability (Kunze et al. 2025).

In response to pulse disturbances, the stabilizing influence of interaction release fades over time. Just after the disturbance, interaction release is the dominant force shaping species responses, enabling compensatory dynamics to emerge. But as the community gradually recovers, this effect diminishes. Eventually, self-regulation takes over as the main driver of stability.

Consider a scenario where a fixed fraction of biomass is removed from all species. In the short term, suppressed species benefit from reduced antagonistic pressures, allowing rapid recovery. Facilitated species, on the other hand, suffer the loss of positive interactions, which slows their rebound. As species regain their biomass, both antagonistic and facilitative interactions return, ending the period of interaction release. At this point, recovery rates are governed by self-regulation.

Previous studies have noted a disconnect between the short- and long-term phases of community recovery (Neubert et al. 1997; Tang et al. 2014; Yang et al. 2023a). These phases are often described using two metrics: reactivity, which captures the short-term response to disturbance, and resilience, which measures the long-term return to equilibrium. Theory suggests that abundant species govern short-term dynamics, while rare species contribute to the long-term recovery (Arnoldi et al. 2018).

We build on these insights by showing that this shift is driven by a transition between two recovery mechanisms. In the short term, species are influenced by interaction release, as disturbances weaken interspecific interactions. As species recover, interactions are restored and the influence of interaction release fades. In the long-term, recovery is governed by self-regulation.

Self-regulation is a key mechanism governing species and community stability. Self-regulation is essential for stable species coexistence (Chesson 2000; May 1972). Dynamical models often require high levels of self-regulation to generate stable yet diverse communities, as observed in nature (Barabás et al. 2017; de Ruiter et al. 1995). However, our results indicate that at the species level, stronger self-regulation does not always equate to greater stability. In particular, species with high SL benefit less from antagonistic release and can even suffer from facilitative release, which hampers their stability when SL ≥ 1, as their high biomass is maintained by their biotic environment. This highlights the nuanced role of self-regulation in shaping species stability.

Our work bridges previously disconnected facets of ecological stability. Stability is indeed defined and measured in many ways (Grimm et al. 1997; Kéfi et al. 2019; Pennekamp et al. 2018a). While previous studies have attempted to link different stability measures (Donohue et al. 2013; Hillebrand et al. 2018; Domínguez-García et al. 2019; Radchuk et al. 2019), they have largely focused on community-level stability and relied on correlative approaches, offering limited mechanistic insights. In contrast, our approach connects different dimensions of stability through a mechanistic understanding of individual species responses. We found that species sensitivity to communitywide press disturbances and short-term recovery from pulse disturbances are both governed by interaction release, while species sensitivity to targeted press disturbances and longterm recovery are determined by self-regulation. Because these pairs of metrics are controlled by the same underlying mechanisms, they exhibit similar patterns and high correlations. In this way, our theory provides a robust foundation for understanding the mechanisms linking seemingly unrelated stability metrics.

Our work has focused on species-level stability to hone a better understanding of the mechanisms that play out. However, many studies have investigated community-level stability (Kéfi et al. 2019). This prompts the question of what SL can tell or not at this higher level of organization. We want to emphasize the difficulty, and therefore the interest, of this question as stability patterns at the species level tend to vanish when aggregated at the community level (SI Section S2; Ives et al. 1999; Loreau et al. 2013).

Furthermore, recent studies have explored how biotic interactions shape species stability under non-equilibrium dynamics (Medeiros et al. 2023; Medeiros et al. 2025). These offer promising directions to extend our framework beyond equilibrium settings. In particular, we find that species SL is closely related to a metric proposed by Medeiros et al. (2023) to predict species sensitivity in dynamic, non-equilibrium contexts (SI Section S8). This correspondence suggests that the mechanistic insights underlying SL may remain informative even when communities are not at steady state.

That said, our framework comes with a key limitation: SL relies on knowing species biomass when grown in isolation. In many cases, such measurements are impractical or impossible. Moreover, some species—such as plants that require facilitation in harsh environments or consumers dependent on their prey—cannot persist alone. We show that SL can be extended to account for these obligate species (SI Section S1). Preliminary investigations suggest that this extended SL remains predictive of species stability (SI, Section S3). However, estimating SL in these cases requires quantifying self-regulation strength, a parameter that is difficult to measure and rarely reported. Developing experimental designs and modeling approaches to address this limitation is an important next step.

The simplicity of the stability patterns we report stems in part from the linearity of the generalized Lotka–Volterra (gLV) model. More generally, the concept of SL separates into two distinct quantities: a timescale ratio that governs response to pulse disturbances, and a biomass ratio that governs response to press disturbances. Despite these nuances, species stability remains clearly structured even in more complex models (SI Section S1).

The breakdown of our theoretical predictions for longterm recovery under pulse disturbances, when species differ in growth rates, highlights a promising direction for future research (SI Section S9). In this case, biotic interactions and species traits—such as generation time or metabolic rate—become entangled, jointly shaping recovery. This opens the door to exploring how our interaction-based framework can be complemented by trait-based approaches. In particular, response diversity—the variation in how species respond to the same disturbance due to differences in life history traits—may help explain patterns not captured by SL alone (Ross et al. 2023; Elmqvist et al. 2003). Integrating these perspectives could yield a more complete picture of species stability in complex ecological communities.

We have shown that SL is a powerful indicator of species stability in systems described by ordinary differential equations. However, not all ecological dynamics fit neatly into this framework. In spatially structured communities, for example, partial differential equations may offer a better description by capturing dispersal across space (Baron et al. 2020). In other cases, time delays—like lags in growth or interaction effects—are better captured with delay differential equations (Yang et al. 2023b). Understanding whether SL remains a useful predictor in these more complex settings is a key next step.

## Acknowledgments

We thank all Biodicée members, P. Kamal, J. Orr and the anonymous reviewer for their valuable suggestions. I.L. and S.K. were supported by the grant ANR-18-CE02-0010-01 of the French National Research Agency ANR (project EcoNet). J-F.A. and M.L. were supported by the ‘Laboratoires d’Excellences (LABEX)’ TULIP (ANR-10-LABX-41).

## Code and data availability

The code used to produce the figures is available on Zenodo at (Lajaaiti 2025). The original data analyzed in this study is available on Zenodo at (Pennekamp et al. 2018b).

## Notes

https://doi.org/10.5281/zenodo.15076241

### Competing Interest Statement

The authors have declared no competing interest.

### Summary of Updates

Figure 1 revised; Figure 4 revised; Methods section moved; Results section revised; Supplemental files updated.

## References

[1] Abrams, Peter A. (June 1983). “Arguments in Favor of Higher Order Interactions”. In: The American Naturalist 121.6, pp. 887–891. ISSN: 0003-0147. DOI: 10.1086/284111.

[2] Abrams, Peter A. et al. (1996). “The Role of Indirect Effects in Food Webs”. In: Food Webs: Integration of Patterns & Dynamics. Ed. by Gary A. Polis and Kirk O. Winemiller. Boston, MA: Springer US, pp. 371–395. ISBN: 978-1-4615-7007-3. DOI: 10.1007/978-1-4615-7007-3_36.

[3] Allesina, Stefano and Si Tang (Mar. 2012). “Stability Criteria for Complex Ecosystems”. In: Nature 483.7388, pp. 205–208. ISSN: 1476-4687. DOI: 10.1038/nature10832.

[4] Allesina, Stefano et al. (July 2015). “Predicting the Stability of Large Structured Food Webs”. In: Nature Communications 6.1, p. 7842. ISSN: 2041-1723. DOI: 10.1038/ncomms8842.

[5] Arnoldi, J. -F. et al. (Jan. 2018). “How Ecosystems Recover from Pulse Perturbations: A Theory of Short-to Long-Term Responses”. In: Journal of Theoretical Biology 436, pp. 79–92. ISSN: 0022-5193. DOI: 10.1016/j.jtbi.2017.10.003.

[6] Barabás, György, Matthew J. Michalska-Smith, and Stefano Allesina (Dec. 2017). “Self-Regulation and the Stability of Large Ecological Networks”. In: Nature Ecology & Evolution 1.12, pp. 1870–1875. ISSN: 2397-334X. DOI: 10.1038/s41559-017-0357-6.

[7] Barbier, Matthieu et al. (Feb. 2018). “Generic Assembly Patterns in Complex Ecological Communities”. In: Proceedings of the National Academy of Sciences 115.9, pp. 2156–2161. ISSN: 0027-8424, 1091-6490. DOI: 10.1073/pnas.1710352115.

[8] Baron, Joseph W. and Tobias Galla (Nov. 2020). “Dispersal-Induced Instability in Complex Ecosystems”. In: Nature Communications 11.1, p. 6032. ISSN: 2041-1723. DOI: 10.1038/s41467-020-19824-4.

[9] Battiston, Federico et al. (Oct. 2021). “The Physics of Higher-Order Interactions in Complex Systems”. In: Nature Physics 17.10, pp. 1093–1098. ISSN: 1745-2481. DOI: 10.1038/s41567-021-01371-4.

[10] Bello, Francesco de et al. (Sept. 2021). “Functional Trait Effects on Ecosystem Stability: Assembling the Jigsaw Puzzle”. In: Trends in Ecology & Evolution 36.9, pp. 822–836. ISSN: 0169-5347. DOI: 10.1016/j.tree.2021.05.001.

[11] Bender, Edward A., Ted J. Case, and Michael E. Gilpin (1984). “Perturbation Experiments in Community Ecology: Theory and Practice”. In: Ecology 65.1, pp. 1–13. ISSN: 1939-9170. DOI: 10.2307/1939452.

[12] Bezanson, Jeff et al. (Sept. 2012). Julia: A Fast Dynamic Language for Technical Computing. DOI: 10.48550/arXiv.1209.5145. arXiv: 1209.5145 [cs].

[13] Bunin, Guy (Apr. 2017). “Ecological Communities with Lotka-Volterra Dynamics”. In: Physical Review E 95.4, p. 042414. DOI: 10.1103/PhysRevE.95.042414.

[14] Capdevila, Pol et al. (2022). “Life History Mediates the Trade-Offs among Different Components of Demographic Resilience”. In: Ecology Letters 25.6, pp. 1566–1579. ISSN: 1461-0248. DOI: 10.1111/ele.14004.

[15] Carrara, Francesco et al. (2015). “Inferring Species Interactions in Ecological Communities: A Comparison of Methods at Different Levels of Complexity”. In: Methods in Ecology and Evolution 6.8, pp. 895–906. ISSN: 2041-210X. DOI: 10.1111/2041-210X.12363.

[16] Chesson, Peter (Nov. 2000). “Mechanisms of Maintenance of Species Diversity”. In: Annual Review of Ecology, Evolution, and Systematics 31.Volume 31, 2000, pp. 343–366. ISSN: 1543-592X, 1545-2069. DOI: 10.1146/annurev.ecolsys.31.1.343.

[17] de Ruiter, Peter C., Anje-Margriet Neutel, and John C. Moore (Sept. 1995). “Energetics, Patterns of Interaction Strengths, and Stability in Real Ecosystems”. In: Science 269.5228, pp. 1257–1260. DOI: 10.1126/science.269.5228.1257.

[18] Domínguez-García, Virginia, Vasilis Dakos, and Sonia Kéfi (Dec. 2019). “Unveiling Dimensions of Stability in Complex Ecological Networks”. In: Proceedings of the National Academy of Sciences 116.51, pp. 25714–25720. DOI: 10.1073/pnas.1904470116.

[19] Donohue, Ian et al. (2013). “On the Dimensionality of Ecological Stability”. In: Ecology Letters 16.4, pp. 421–429. ISSN: 1461-0248. DOI: 10.1111/ele.12086.

[20] Elmqvist, Thomas et al. (2003). “Response Diversity, Ecosystem Change, and Resilience”. In: Frontiers in Ecology and the Environment 1.9, pp. 488–494. ISSN: 1540-9309. DOI: 10.1890/1540-9295(2003)001[0488:RDECAR]2.0.CO;2.

[21] Gamelon, Marlène et al. (Nov. 2014). “Influence of Life-History Tactics on Transient Dynamics: A Comparative Analysis across Mammalian Populations.” In: The American Naturalist 184.5, pp. 673–683. ISSN: 0003-0147. DOI: 10.1086/677929.

[22] Gonzalez, Andrew and Michel Loreau (Dec. 2009). “The Causes and Consequences of Compensatory Dynamics in Ecological Communities”. In: Annual Review of Ecology, Evolution, and Systematics 40.Volume 40, 2009, pp. 393–414. ISSN: 1543-592X, 1545-2069. DOI: 10.1146/annurev.ecolsys.39.110707.173349.

[23] Grimm, Volker and Christian Wissel (1997). “Babel, or the Ecological Stability Discussions: An Inventory and Analysis of Terminology and a Guide for Avoiding Confusion”. In: Oecologia 109.3, pp. 323–334. ISSN: 0029-8549. JSTOR: 4221528.

[24] Grman, Emily et al. (2010). “Mechanisms Contributing to Stability in Ecosystem Function Depend on the Environmental Context”. In: Ecology Letters 13.11, pp. 1400–1410. ISSN: 1461-0248. DOI: 10.1111/j.1461-0248.2010.01533.x.

[25] Guimarães, Paulo R. (Nov. 2020). “The Structure of Ecological Networks Across Levels of Organization”. In: Annual Review of Ecology, Evolution, and Systematics 51.Volume 51, 2020, pp. 433–460. ISSN: 1543-592X, 1545-2069. DOI: 10.1146/annurev-ecolsys-012220-120819.

[26] Hillebrand, Helmut et al. (2018). “Decomposing Multiple Dimensions of Stability in Global Change Experiments”. In: Ecology Letters 21.1, pp. 21–30. ISSN: 1461-0248. DOI: 10.1111/ele.12867.

[27] Ives, A. R., K. Gross, and J. L. Klug (Oct. 1999). “Stability and Variability in Competitive Communities”. In: Science 286.5439, pp. 542–544. DOI: 10.1126/science.286.5439.542.

[28] Kardol, Paul et al. (2010). “Climate Change Effects on Plant Biomass Alter Dominance Patterns and Community Evenness in an Experimental Old-Field Ecosystem”. In: Global Change Biology 16.10, pp. 2676–2687. ISSN: 1365-2486. DOI: 10.1111/j.1365-2486.2010.02162.x.

[29] Kawatsu, Kazutaka (July 2024). “Unraveling Emergent Network Indeterminacy in Complex Ecosystems: A Random Matrix Approach”. In: Proceedings of the National Academy of Sciences 121.27, e2322939121. DOI: 10.1073/pnas.2322939121.

[30] Kéfi, Sonia et al. (2012). “More than a Meal… Integrating Non-Feeding Interactions into Food Webs”. In: Ecology Letters 15.4, pp. 291–300. ISSN: 1461-0248. DOI: 10.1111/j.1461-0248.2011.01732.x.

[31] Kéfi, Sonia et al. (2019). “Advancing Our Understanding of Ecological Stability”. In: Ecology Letters 22.9, pp. 1349–1356. ISSN: 1461-0248. DOI: 10.1111/ele.13340.

[32] Kunze, Charlotte et al. (2025). “Partitioning Species Contributions to Ecological Stability in Disturbed Communities”. In: Ecological Monographs 95.1, e1636. ISSN: 1557-7015. DOI: 10.1002/ecm.1636.

[33] Lajaaiti, Ismaël (June 2025). Ismael-Lajaaiti/Species-Stability: Submission for ELE. Zenodo. DOI: 10.5281/zenodo.15632312.

[34] Lajaaiti, Ismaël, Sonia Kéfi, and Jean-François Arnoldi (Oct. 2024). “How Biotic Interactions Structure Species’ Responses to Perturbations”. In: Proceedings of the Royal Society B: Biological Sciences 291.2032, p. 20240930. DOI: 10.1098/rspb.2024.0930.

[35] Lavorel, S. and E. Garnier (2002). “Predicting Changes in Community Composition and Ecosystem Functioning from Plant Traits: Revisiting the Holy Grail”. In: Functional Ecology 16.5, pp. 545–556. ISSN: 1365-2435. DOI: 10.1046/j.1365-2435.2002.00664.x.

[36] Loreau, M. et al. (Oct. 2001a). “Biodiversity and Ecosystem Functioning: Current Knowledge and Future Challenges”. In: Science 294.5543, pp. 804–808. DOI: 10.1126/science.1064088.

[37] Loreau, Michel and Claire de Mazancourt (2013). “Biodiversity and Ecosystem Stability: A Synthesis of Underlying Mechanisms”. In: Ecology Letters 16.1, pp. 106–115. ISSN: 1461-0248. DOI: 10.1111/ele.12073.

[38] Loreau, Michel and Andy Hector (July 2001b). “Partitioning Selection and Complementarity in Biodiversity Experiments”. In: Nature 412.6842, pp. 72–76. ISSN: 1476-4687. DOI: 10.1038/35083573.

[39] Lubiana Botelho, Larissa et al. (2025). “Calibrated Ecosystem Models Cannot Predict the Consequences of Conservation Management Decisions”. In: Ecology Letters 28.1, e70034. ISSN: 1461-0248. DOI: 10.1111/ele.70034.

[40] Mariotte, Pierre et al. (2013a). “Subordinate Plant Species Enhance Community Resistance against Drought in Semi-Natural Grasslands”. In: Journal of Ecology 101.3, pp. 763–773. ISSN: 1365-2745. DOI: 10.1111/1365-2745.12064.

[41] Mariotte, Pierre et al. (Apr. 2013b). “Subordinate Plant Species Impact on Soil Microbial Communities and Ecosystem Functioning in Grasslands: Findings from a Removal Experiment”. In: Perspectives in Plant Ecology, Evolution and Systematics 15.2, pp. 77–85. ISSN: 1433-8319. DOI: 10.1016/j.ppees.2012.12.003.

[42] May, Robert M. (Aug. 1972). “Will a Large Complex System Be Stable?” In: Nature 238.5364, pp. 413–414. ISSN: 1476-4687. DOI: 10.1038/238413a0.

[43] May, Robert M. (Dec. 2019). Stability and Complexity in Model Ecosystems. Princeton University Press. ISBN: 978-0-691-20691-2.

[44] McDonald, Jenni L. et al. (Jan. 2017). “Divergent Demographic Strategies of Plants in Variable Environments”. In: Nature Ecology & Evolution 1.2, pp. 1–6. ISSN: 2397-334X. DOI: 10.1038/s41559-016-0029.

[45] Medeiros, Lucas P. et al. (2023). “Ranking Species Based on Sensitivity to Perturbations under Non-Equilibrium Community Dynamics”. In: Ecology Letters 26.1, pp. 170–183. ISSN: 1461-0248. DOI: 10.1111/ele.14131.

[46] Medeiros, Lucas P. et al. (May 2025). A Nonequilibrium Framework for Community Responses to Pulse Perturbations. DOI: 10.1101/2025.05.14.654148.

[47] Mentges, Andrea et al. (2024). “Accounting for Effects of Growth Rate When Measuring Ecological Stability in Response to Pulse Perturbations”. In: Ecology and Evolution 14.10, e11637. ISSN: 2045-7758. DOI: 10.1002/ece3.11637.

[48] Montoya, José M., Stuart L. Pimm, and Ricard V. Solé (July 2006). “Ecological Networks and Their Fragility”. In: Nature 442.7100, pp. 259–264. ISSN: 1476-4687. DOI: 10.1038/nature04927.

[49] Neubert, Michael G. and Hal Caswell (1997). “Alternatives to Resilience for Measuring the Responses of Ecological Systems to Perturbations”. In: Ecology 78.3, pp. 653–665. ISSN: 1939-9170. DOI: 10.1890/0012-9658(1997)078[0653:ATRFMT]2.0.CO;2.

[50] Novak, Mark et al. (Nov. 2016). “Characterizing Species Interactions to Understand Press Perturbations: What Is the Community Matrix?” In: Annual Review of Ecology, Evolution, and Systematics 47.Volume 47, 2016, pp. 409–432. ISSN: 1543-592X, 1545-2069. DOI: 10.1146/annurev-ecolsys-032416-010215.

[51] Patten, Bernard C. (Feb. 1982). “Environs: Relativistic Elementary Particles for Ecology”. In: The American Naturalist 119. ISSN: 0003-0147. DOI: 10.1086/283903.

[52] Pennekamp, Frank et al. (Nov. 2018a). “Biodiversity Increases and Decreases Ecosystem Stability”. In: Nature 563.7729, pp. 109–112. ISSN: 1476-4687. DOI: 10.1038/s41586-018-0627-8.

[53] Pennekamp, Frank et al. (Aug. 2018b). Pennekampster/Code_and_data_OverallEcosystemStability: Release of Data and Code. Zenodo. DOI: 10.5281/zenodo.1345557.

[54] Pichon, Benoît et al. (2024). “Integrating Ecological Feed-backs across Scales and Levels of Organization”. In: Ecography n/a.n/a, e07167. ISSN: 1600-0587. DOI: 10.1111/ecog.07167.

[55] Rackauckas, Christopher and Qing Nie (May 2017). “DifferentialEquations.Jl – A Performant and Feature-Rich Ecosystem for Solving Differential Equations in Julia”. In: Journal of Open Research Software 5.1, pp. 15–15. ISSN: 2049-9647. DOI: 10.5334/jors.151.

[56] Radchuk, Viktoriia et al. (2019). “The Dimensionality of Stability Depends on Disturbance Type”. In: Ecology Letters 22.4, pp. 674–684. ISSN: 1461-0248. DOI: 10.1111/ele.13226.

[57] Ross, Samuel R. P.-J. et al. (2023). “How to Measure Response Diversity”. In: Methods in Ecology and Evolution 14.5, pp. 1150–1167. ISSN: 2041-210X. DOI: 10.1111/2041-210X.14087.

[58] Tang, Si and Stefano Allesina (2014). “Reactivity and Stability of Large Ecosystems”. In: Frontiers in Ecology and Evolution 2. ISSN: 2296-701X.

[59] Terry, J. Christopher D. (Apr. 2025). Unfeasible Expectations: Why Simple Predictors Outperform Structural Stability Measures for Understanding Community Assembly. DOI: 10.1101/2025.04.24.650476.

[60] Yang, Yuguang et al. (Nov. 2023a). “Reactivity of Complex Communities Can Be More Important than Stability”. In: Nature Communications 14.1, p. 7204. ISSN: 2041-1723. DOI: 10.1038/s41467-023-42580-0.

[61] Yang, Yuguang et al. (Oct. 2023b). “Time Delays Modulate the Stability of Complex Ecosystems”. In: Nature Ecology & Evolution 7.10, pp. 1610–1619. ISSN: 2397-334X. DOI: 10.1038/s41559-023-02158-x.

[62] Yodzis, Peter (1988). “The Indeterminacy of Ecological Interactions as Perceived Through Perturbation Experiments”. In: Ecology 69.2, pp. 508–515. ISSN: 1939-9170. DOI: 10.2307/1940449.

[63] Zelnik, Yuval R. et al. (2024). “How Collectively Integrated Are Ecological Communities?” In: Ecology Letters 27.1, e14358. ISSN: 1461-0248. DOI: 10.1111/ele.14358.

