## Supporting Information for "Linking biotic interactions to species stability"

### SUPPORTING INFORMATION TO
### Linking biotic interactions to species stability

Ismaël Lajaaiti[1,*], Sonia Kéfi[1,2], Michel Loreau [3,4],
Alice Ardichvili[3] and Jean-François Arnoldi[3]

[1]ISEM, CNRS, Univ. de Montpellier, IRD, Montpellier, France
[2]Santa Fe Institute, 1399 Hyde Park Road, Santa Fe, NM 87501, USA
[3]Theoretical and Experimental Ecology Station, CNRS, 2 route du CNRS, 09200, Moulis, France
[4]Institute of Ecology, College of Urban and Environmental Sciences, Peking University, Beijing 100871, China

### S1 Generalization to non-linear models

In the main text, we highlight the close overlap between self-regulation loss (SL) and relative yield (RY). This similarity arises from the linearity of the generalized Lotka-Volterra (gLV) model used in our study, where SL and RY are mathematically equivalent:

$$\mathrm{SL}_i = \mathrm{RY}_i = \frac{B_i}{K_i}$$

where $B_i$ is the species biomass in a community, and $K_i$ its carrying capacity. As a result, both metrics may appear to capture the organization of species stability. Here, we demonstrate that this equivalence vanishes in non-linear models, allowing SL and RY to be disentangled. More importantly, we show that species charateristic recovery time scale is governed by SL, while the effect of antagonistic of facilitation release is determined by a measure related to RY, that captures net effects of biotic interactions.

## SL v RY

We consider a more general class of species dynamics, given by

$$\frac{\mathrm{d}B_i}{\mathrm{d}t} = B_i f_i(\mathbf{B})$$

where $B_i$ is the biomass of species $i$, $\mathbf{B}$ is the vector of species biomasses, and $f_i$ is the per capita growth rate of species $i$, which can be a linear or non-linear function of species biomass. In this framework, we define species SL as

$$\mathrm{SL}_i = \frac{-\frac{\partial f_i}{\partial B_i} B_i}{f_i(0)}.$$

This expression represents the proportion of biomass loss due to self-regulation, normalized by the species' intrinsic growth rate at zero biomass. The normalization by $f_i(0)$ ensures fair comparison between species with different growth rates. In the case of the gLV model, we recover

$$f_i(\mathbf{B}) = r_i \left(1 + \frac{-B_i + \sum_{j \neq i} a_{ij} B_j}{K_i}\right), \quad \mathrm{SL}_i = \frac{B_i}{K_i}.$$

On the other hand, the traditional definition of RY (Loreau and Hector 2001) remains

$$\mathrm{RY}_i = \frac{B_i}{K_i}.$$

To further distinguish SL from RY, we explicitly derive the expression for species carrying capacity in the general case:

$$K_i = -\frac{f_i(0)}{\langle \frac{\partial f_i}{\partial B_i} \rangle}, \quad \text{where} \quad \langle \frac{\partial f_i}{\partial B_i} \rangle = \frac{1}{K_i} \int_0^{K_i} \frac{\partial f_i}{\partial B_i} \mathrm{d}B_i.$$

This formula can be understood graphically by looking at Fig. S1A. The average slope of species per capita growth is the ratio of its intrinsic growth rate to its carrying capacity. Using this expression, we can write RY as

$$\mathrm{RY}_i = \frac{-\langle \frac{\partial f_i}{\partial B_i} \rangle B_i}{f_i(0)}.$$

This formulation highlights a key difference: SL depends on the *local* derivative of the species' per capita growth rate, whereas RY depends on its *average* derivative (Fig. S1A). In the gLV model, where $f_i$ is linear, these values are equal ($-\frac{r_i}{K_i}$), but this is not generally the case (Fig. S1B).

#### $\theta$-logistic growth

To illustrate the difference between SL and RY, we consider the the $\theta$-logistic growth model (Sibly et al. 2005), that generalizes the logistic growth. In this model, species growth is given by

$$f_i(B_i) = \frac{r}{\theta}\left[1 - \left(\frac{B_i}{K_i}\right)^{\theta}\right].$$

When $\theta = 1$, we recover the logistic growth. When $-1 < \theta < 0$, species growth is sub-linear in its biomass (**hatton2024diversity**). We plot this expression for different $\theta$ values in Fig. S1B. We can compute species SL in this model and we find that

$$\mathrm{SL}_i = \mathrm{RY}_i^{\theta}.$$

Notably, this formula implies that for negative $\theta$ values, SL and RY should be negatively correlated, as observed in Fig. S1C.

Having established that SL and RY are distinct, we will now demonstrate that: 1) species characteristic recovery rate is determined by SL, and 2) species response to antagonistic release is determined by a term close to RY.

#### Characteristic recovery rate

Including interspecific interactions species growth is governed by

$$\frac{1}{B_i}\frac{\mathrm{d}B_i}{\mathrm{d}t} = \frac{r}{\theta}\left[1 - \left(\frac{B_i}{K_i}\right)^{\theta}\right] + \frac{\sum_{j\neq i} A_{ij} B_j}{K_i}.$$

We recover the Lotka-Volterra equation presented in the main text for $\theta = 1$, when species follow a logistic growth. We consider the linearized dynamics of species deviation near their equilibrium $x_i = B_i - B_i^{\mathrm{eq}}$

$$\frac{\mathrm{d}x_i}{\mathrm{d}t} = B_i^{\mathrm{eq}}(A_{ii}x_i + \sum_{j\neq i} A_{ij}x_j), \text{ where } A_{ij} = \frac{\partial f_i}{\partial B_j}.$$

Because we want to compare fairly species with different equilibrium biomass we focus on the recovery of the proportion of biomass lost $z_i = \frac{x_i}{B_i^{\mathrm{eq}}}$, which yields

$$\frac{\mathrm{d}z_i}{\mathrm{d}t} = A_{ii}B_i^{\mathrm{eq}}(z_i - \frac{1}{B_i^{\mathrm{eq}}}\sum_{j\neq i} a_{ij}B_j^{\mathrm{eq}}z_j), \text{ with } a_{ij} = -\frac{A_{ij}}{A_{ii}}.$$

Moreover because we want to fairly compare the recovery of slow and fast species, we divide their growth by $f_i(0)$ which captures the intrinsic population speed (e.g. generation time) (Mentges et al. 2024). This value for instance corresponds to $r_i$ in the gLV model. We obtain

$$\frac{1}{f_i(0)}\frac{\mathrm{d}z_i}{\mathrm{d}t} = -\mathrm{SL}_i(z_i - \frac{1}{B_i^{\mathrm{eq}}}\sum_{j\neq i} a_{ij}B_j^{\mathrm{eq}} z_j).$$

We observe in the equation above that species recovery is globally governed by their SL, demonstrating that this is the relevant measure to capture the species propensity to return to its equilibrium.

In particular, if we ignore interspecific interactions we obtain the expression of species characteristic recovery

$$\frac{1}{f_i(0)}\frac{\mathrm{d}}{\mathrm{d}t}\log z_i = -\mathrm{SL}_i$$

Therefore, species characteristic recovery equates their SL, not RY. We illustrate this point in simulated communities using the $\theta$-logistic model in Fig. S1D,E.

#### Interaction release in response to pulse

We have ignored above the effect of interspecific interactions on species stability (apart from their influence on species equilibrium biomass), however we know that interactions are expected to shape species responses to disturbances. We address this question in this section.

We can do a similar approximation that we have done in the main text: assuming that $a_{ij}B_j^{\mathrm{eq}}$ are weakly correlated to $z_j$, which yields

$$\frac{1}{f_i(0)}\frac{\mathrm{d}z_i}{\mathrm{d}t} = -\mathrm{SL}_i(z_i - \langle z\rangle \frac{1}{B_i^{\mathrm{eq}}}\sum_{j\neq i} a_{ij}B_j^{\mathrm{eq}}), \text{ where } \langle z\rangle = \frac{1}{S-1}\sum_{j\neq i} z_j \approx \frac{1}{S}\sum_j z_j.$$

We define the pseudo-carrying capacity as

$$\hat{K}_i = B_i^{\mathrm{eq}} - \sum_{j\neq i} a_{ij}B_j^{\mathrm{eq}}.$$

In the gLV model this sum equates species carrying capacity $K_i$, yet, in nonlinear model we have $K_i \neq \hat{K}_i$. Next, we can naturally define a pseudo relative yield

$$\hat{\mathrm{RY}}_i = \frac{B_i^{\mathrm{eq}}}{\hat{K}_i}.$$

When competition is strong locally, that is near the equilibrium ($a_{ij}$ can be density dependent), we have that $\hat{K}_i \ll B_i^{\mathrm{eq}}$ and therefore $\hat{\mathrm{RY}}_i \ll 1$. By contrast, when facilitation is strong, we have $\hat{K}_i \gg B_i^{\mathrm{eq}}$ and therefore $\hat{\mathrm{RY}}_i \gg 1$. As a result, the intuitions from the original RY concept still holds, provided that one consider interactions locally. Yet, it appears that RY approximates well $\hat{\mathrm{RY}}$ that the range of $\theta$ considered (Fig. S1F). However, we warn that this may not be the case in all generality.

That being said, we next simplify our equations

$$\frac{1}{f_i(0)}\frac{\mathrm{d}z_i}{\mathrm{d}t} = -\mathrm{SL}_i\Big(z_i + \langle z\rangle(\frac{1}{\hat{\mathrm{RY}}} - 1)\Big).$$

We find that the strength of interaction release is captured by $\frac{1}{\hat{\mathrm{RY}}_i} - 1$, while SL reflects the species' timescale. When species experience strong antagonistic interactions ($\hat{\mathrm{RY}}_i \ll 1$), this term becomes large and positive, highlighting the stabilizing effect of antagonistic release. Conversely, when a species benefits strongly from facilitative interactions ($\hat{\mathrm{RY}}_i \gg 1$), the term becomes negative, indicating the destabilizing effect of facilitative release.

What differs from the gLV model is that, $\mathrm{SL} \neq \mathrm{RY}$, and therefore interaction release and self-regulation are in general decoupled. Specifically, in the gLV the stabilizing effect of interaction release declines with species self-regulation as $\frac{1}{\mathrm{SL}} - 1$, resulting in a balance of the two stabilizing mechanisms which vanishes when considering a broader class of nonlinear model.

We have shown in the main text that interaction release predominantly governs species short-term recovery following the pulse disturbance.The return rate at short-term can be approximated as

$$R_{\mathrm{short},i} = -\frac{1}{f_i(0)}\frac{\mathrm{d}}{\mathrm{d}t}\log z_i \simeq \mathrm{SL}_i\left[1 + \langle z_0\rangle\langle\frac{1}{z_0}\rangle(\frac{1}{\hat{\mathrm{RY}}} - 1)\right]$$

where $z_0$ corresponds to the initial perturbation, which we assume follows the same distribution for all species. $\langle\cdot\rangle$ denotes the average on many disturbance events. We plot this prediction in Fig. S1G, replacing $\hat{\mathrm{RY}}$ by RY.

#### Interaction release in response to press

Having demonstrated how interaction release operate in pulse disturbances, we can now consider the case of press disturbances. Let's consider a small, sustained change in species per capita growth rate. The resulting change in its equilibrium biomass writes

$$\delta \mathbf{B}^{\mathrm{eq}} = (D(\mathbb{I} - A))^{-1}\delta \mathbf{f}, \text{ where } D = \mathrm{Diag}\Big((A_{ii})_{1\leq i \leq S}\Big) \text{ and } (A)_{ij} = \begin{cases} a_{ij} = \frac{A_{ij}}{A_{ii}} & \text{if } i \neq j \\ 0 & \text{otherwise.} \end{cases}$$

We can re-write this as

$$\delta \mathbf{B}^{\mathrm{eq}} = (\mathbb{I} - A)^{-1} D^{-1}\delta \mathbf{f}$$

to define $V = (\mathbb{I} - A)^{-1}$ the sensitivity matrix, whose elements are dimensionless. We recognize that our definition of pseudo carrying capacity can be written

$$\hat{\mathbf{K}} = (\mathbb{I} - A)\mathbf{B}^{\mathrm{eq}}$$

and therefore

$$\mathbf{B}^{\mathrm{eq}} = (\mathbb{I} - A)^{-1}\hat{\mathbf{K}} = V\hat{\mathbf{K}}. \tag{S1}$$

By consistency with this equation, we re-write the response to the press as

$$\delta \mathbf{B}^{\mathrm{eq}} = V\delta\hat{\mathbf{K}}, \text{ where } \delta\hat{\mathbf{K}} = D^{-1}\delta \mathbf{f}.$$

Then, the sensitivity of species $i$ can be decomposed as

$$\frac{\delta B_i^{\mathrm{eq}}}{B_i^{\mathrm{eq}}} = \frac{V_{ii}}{\hat{\mathrm{RY}}_i}\frac{\delta \hat{K}_i}{\hat{K}_i} + \frac{1}{B_i^{\mathrm{eq}}}\sum_{j\neq i} V_{ij}\hat{K}_j\frac{\delta \hat{K}_j}{\hat{K}_j}.$$

As in the main-text, we do a mean-field approximation to simplify the sum of interspecific net effects

$$\frac{\delta B_i^{\text{eq}}}{B_i^{\text{eq}}} = \frac{V_{ii}}{\hat{\text{RY}}_i}\frac{\delta \hat{K}_i}{\hat{K}_i} + \frac{1}{B_i^{\text{eq}}}\sum_{j\neq i} V_{ij}\hat{K}_j\langle\frac{\delta \hat{K}}{\hat{K}}\rangle \text{ where } \langle\frac{\delta \hat{K}}{\hat{K}}\rangle = \frac{1}{S-1}\sum_{j\neq i}\frac{\delta \hat{K}_j}{\hat{K}_j} \approx \frac{1}{S}\sum_j \frac{\delta \hat{K}_j}{\hat{K}_j}.$$

We use Eq. S1 to simplify the sum of interspecific effects

$$\sum_{j\neq i} V_{ij}\hat{K}_j = B_i^{\text{eq}} - V_{ii}K_{ii}$$

which yields

$$\frac{\delta B_i^{\text{eq}}}{B_i^{\text{eq}}} = \frac{V_{ii}}{\hat{\text{RY}}_i}\frac{\delta \hat{K}_i}{\hat{K}_i} - \left(\frac{V_{ii}}{\hat{\text{RY}}_i} - 1\right)\langle\frac{\delta \hat{K}}{\hat{K}}\rangle.$$

By dividing the biomass change by the press intensity on species $i$, and averaging on many disturbances, we obtain

$$s_i = \frac{V_{ii}}{\hat{\text{RY}}_i} - \langle\kappa\rangle\langle\frac{1}{\kappa}\rangle\left(\frac{V_{ii}}{\hat{\text{RY}}_i} - 1\right), \text{ where } \kappa = \frac{\delta \hat{K}}{\hat{K}}.$$

The effect of interaction release on stability is governed by $\frac{V_{ii}}{\hat{\text{RY}}i} - 1$, closely resembling the expression derived for pulse disturbances. This aligns with our earlier findings: species experiencing strong antagonistic interactions gain stability through antagonistic release, while facilitated species experience reduced stability due to facilitative release. Specifically, when interactions are relatively weak—typically allowing full species coexistence—we observe that $Vii \simeq 1$ (see Fig.S4). We validate this prediction by replacing $\hat{\text{RY}}$ with RY in Fig.S1F.

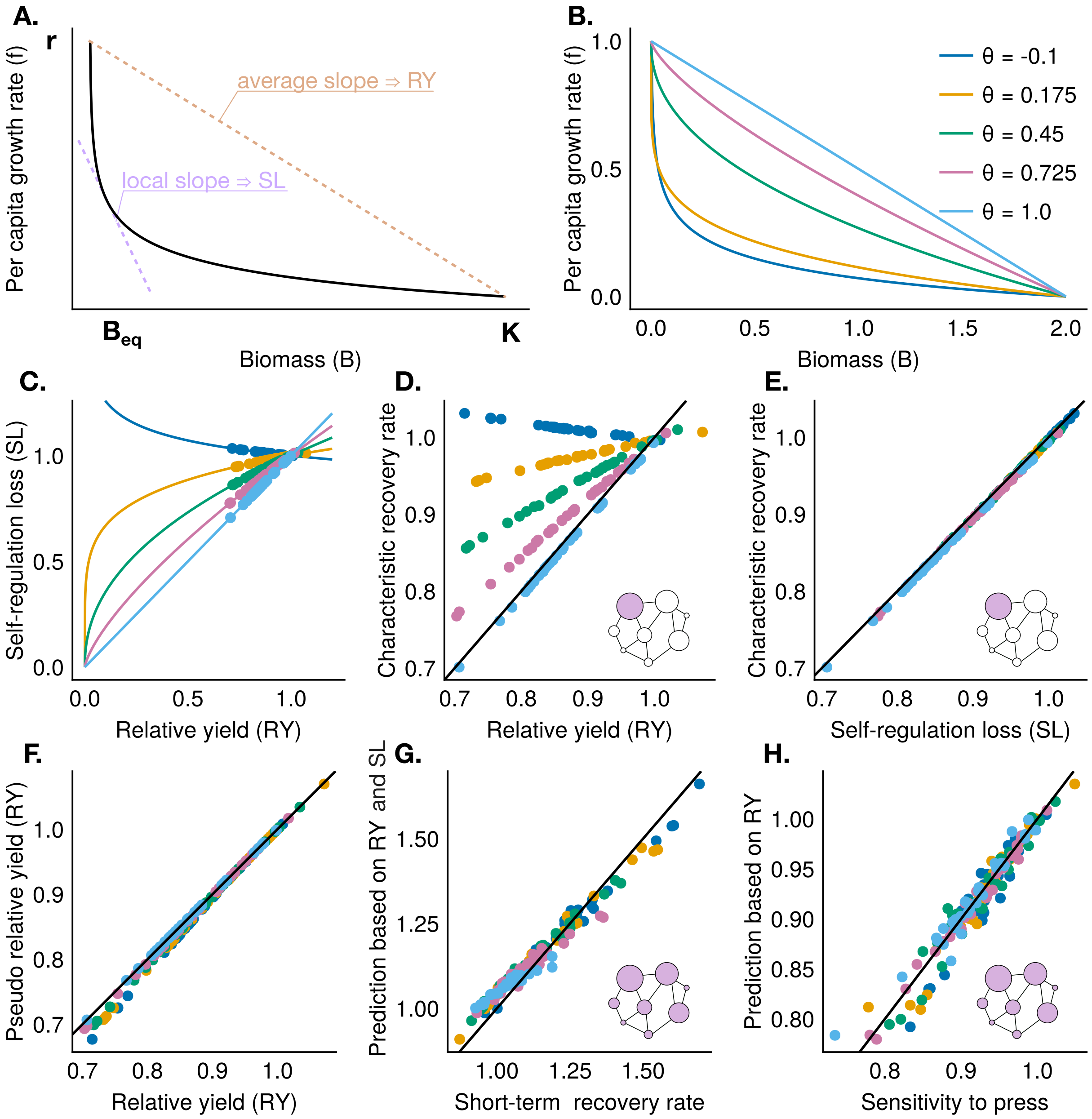


Figure S1: Organization of species stability in the $\theta$-logistic model. (A) Graphical illustration of the dependency of SL on different features of the species per capita growth curve. (B) Species per capita growth under the $\theta$-logistic model for different $\theta$ values. (C) Species SL against their RY, showing that both metrics are not equal and become negatively correlated for sublinear growth ($\theta < 0$). (D) Species relative yield does not capture species characteristic recovery rate beyond the logistic growth ($\theta = 1$). Characteristic recover rate is the species recovery rate following a pulse targeting it (other species undisturbed). Initial disturbance was set to $z_i(0) = -0.01$. Recovery rate was computed between $t = 0$ and $t = 1$. (E) Characteristic recovery rate against SL. (F) Pseudo relative yield against relative yield. (G) Predicted vs. observed values of species short-term recovery rates. Initial disturbances were drawn from $z_i(0) \sim -\text{LogNormal}(\log 0.1, 0.65)$ (all species simultaneously affected). Recovery rates were averaged over 500 disturbance events. (H) Predicted vs. observed values of species sensitivity to community-wide press. The press was model as a decrease in species carrying capacity. Press intensity were drawn from $\frac{\delta K_i}{K_i} \sim -0.1\text{LogNormal}(\log 0.1, 0.65)$. Sensitivity was averaged over 500 disturbance events. Community parameters: $S = 30$, $S\mu = -0.15$, $\sqrt{S}\sigma = 0.1$, $\text{std}(K_i) = 0.3$, $r_i = 1$.

#### S2 From species to community stability

Do stability patterns observed at the species level necessarily scale up to the community level? Studies on temporal variability suggest that they do not. An increase in species-level stability does not necessarily lead to greater stability of the entire community, often measured by the stability of total community biomass. In particular, research has shown that when population variability rises due to increased competition, species fluctuations become more asynchronous. These two effects counterbalance each other, resulting in community biomass variability that remains independent of competition strength (Ives et al. 1999; Loreau and de Mazancourt 2013).

Here, we demonstrate that this principle extends beyond temporal variability to species responses under both pulse (instantaneous biomass removal) and press (sustained environmental change) disturbances. In Fig. S2, we plot average species stability and community stability against the mean interaction strength ($\mu$) in simulated communities. At the species level, increasing competition (i.e., more negative values of $\mu$) enhances stability due to the strengthening of antagonistic release. However, when aggregated at the community level, this stabilizing effect vanishes, leading to a community-wide stability that remains largely independent of interaction strength. These examples highlight the complexity of scaling stability patterns from species to the community level, showing the need for further research to bridge this gap.

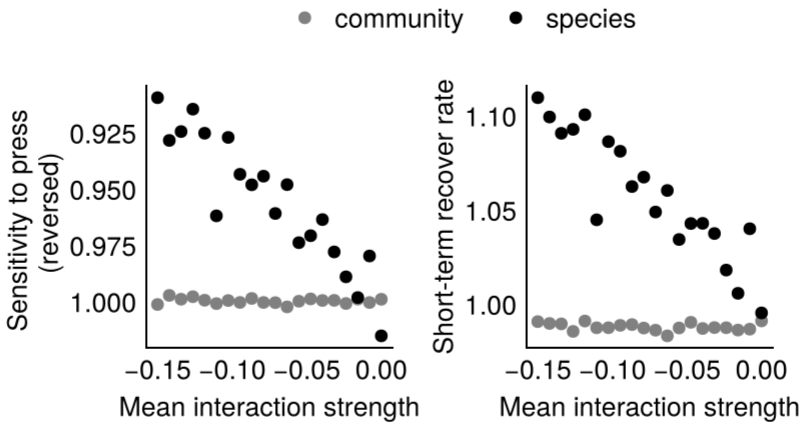


Figure S2: Stability patterns at the species-level vanish at the community-level. (A) Sensitivity to press of species individual biomass (averaged on all species), or of the community total biomass, against mean interaction strength. Press is modeled as a change in species carrying capacity. Press intensities are taken from $\frac{\delta K_i}{K_i} \sim -0.1\text{LogNormal}(\log 0.1, 0.65)$. Sensitivity is averaged on 100 perturbation events. (B) Short-term recovery rate of species individual biomass (averaged on all species), and of community total biomass. Short-term recovery rate is computed between $t = 0$ and $t = 1$. Pulse intensities, $x_i$, are taken from $\frac{\delta x_i}{B_i^{\text{eq}}} \sim -0.1\text{LogNormal}(\log 0.1, 0.65)$. Recovery rate is average on 100 perturbations events. Community parameters: $S = 30$, $S\mu = -0.15$, $\sqrt{S}\sigma = 0.1$, $\text{std}(K_i) = 0.3$, $r_i = 1$.

### S3 Generalization to obligate species

In the main text, we focus on species that can survive in isolation, that is, with $f_i(0) > 0$. Yet, many species depend on others to survive—consumers being a blatant example. We refer to those as dependent species. We explain in this section how our framework extends to such species. For simplicity, we return to the simple, and linear gLV model

$$f_i(\mathbf{B}) = r_i \left( u_i + \frac{-B_i + \sum_{j \neq i} a_{ij} B_j}{K_i} \right).$$

Previously, we had $u_i = 1$ for all species, such that species equilibrium biomass in isolation was given by $K_i > 0$ by definition. We now consider the possibility of negative $u_i$. In such setting, species equilibrium biomass in isolation is given by $u_i K_i < 0$ signaling that species depend on others for their survival.

By extending the notion of RY (or SL), to

$$\mathrm{RY}_i = \frac{B_i}{u_i K_i},$$

we will show that the reasoning presented in the main text holds.

At equilibrium we have

$$\mathbf{uK} = -(\mathbb{I} - A)\mathbf{B}^{\mathrm{eq}}.$$

Then species equilibrium are given by

$$\mathbf{B}^{\mathrm{eq}} = V\mathbf{uK},$$

with $V = -(\mathbb{I} - A)^{-1}$ the sensitivity matrix. We model press disturbance as a shift in $u_i$. The resulting change in species equilibrium biomass writes

$$\Delta\mathbf{B}^{\mathrm{eq}} = V\delta\mathbf{uK},$$

that is for species $i$

$$\Delta B_i^{\mathrm{eq}} = \sum_j V_{ij} \delta u_j K_j.$$

Because, we consider that shift in $u_i$ are drawn proportionally to $u_i$, we write

$$\frac{\Delta B_i^{\mathrm{eq}}}{B_i^{\mathrm{eq}}} = V_{ii} \frac{\delta u_i}{u_i} \frac{u_i K_i}{B_i^{\mathrm{eq}}} + \frac{1}{B_i^{\mathrm{eq}}} \sum_{j \neq i} V_{ij} u_j K_j \frac{\delta u_j}{u_j}.$$

We recognize the generalized relative yield, and we can simplify the rightmost sum,

$$\frac{\Delta B_i^{\mathrm{eq}}}{B_i^{\mathrm{eq}}} = \frac{V_{ii}}{\mathrm{RY}_i} \frac{\delta u_i}{u_i} + \frac{1}{B_i^{\mathrm{eq}}} \sum_{j \neq i} V_{ij} u_j K_j \langle \frac{\delta u}{u} \rangle.$$

Moreover, we have that

$$\frac{1}{B_i^{\mathrm{eq}}} \sum_{j \neq i} V_{ij} u_j K_j = 1 - \frac{V_{ii}}{\mathrm{RY}_i}$$

which yields

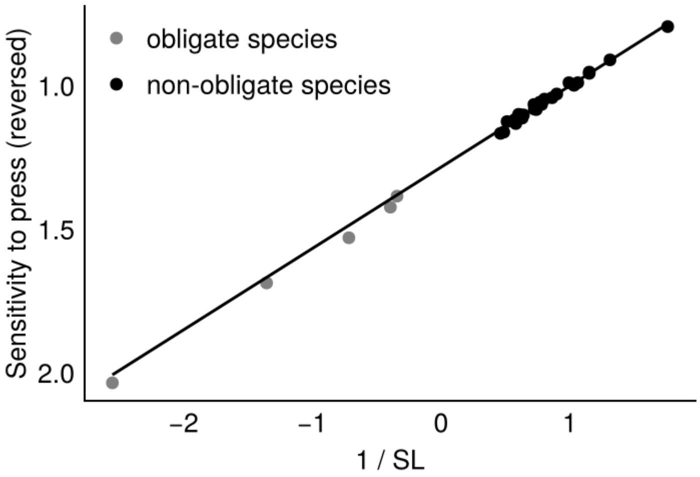


Figure S3: Generalization of our predictions of species sensitivity ($s$) to press to dependent species. We model press disturbances as a shift in species growth $u_i$. Press intensities are drawn in $\delta u_i \sim \text{LogNormal}(-1, 0.5)u_i$. Sensitivity is averaged over 1,000 disturbance events. Community parameters: $S = 30$, $S\mu = 0.3$, $\sqrt{S}\sigma = 0.3$, $\text{std}(K_i) = 0.3$, $r_i = 1$, $u_i = -0.1$ for dependent species ($S_d = 5$) and $u_i = 0.4$ for non-dependent species ($S_{nd} = 25$).

$$\frac{\Delta B_i^{\text{eq}}}{B_i^{\text{eq}}} = \frac{V_{ii}}{\text{RY}_i}\frac{\delta u_i}{u_i} + (1 - \frac{V_{ii}}{\text{RY}_i})\langle\frac{\delta u}{u}\rangle.$$

Dividing by the intensity of the press on species $i$ to obtain the sensitivity, and averaging on many disturbance events give

$$s_i = \frac{V_{ii}}{\text{RY}_i} + (1 - \frac{V_{ii}}{\text{RY}_i})\langle\frac{\delta u}{u}\rangle\langle(\frac{\delta u}{u})^{-1}\rangle.$$

We recognize the formula derived in the main text, provided that we extend the notion of relative yield as explained above. We test this prediction on simulation in Fig. S3, and shows that it accurately predicts species sensitivity, even for species with $u_i < 0$.

#### S4 Community feedback

In this section, we derive an analytical expression of species feedback $V_{ii}$ depending on the interaction parameters.

We consider the introduction of species 0 in the community. For conciseness we write the RY of species, $\eta_i$. Morever, we write the relative yield of species $i$ in absence of species 0, $\eta_{i_{/0}}$. In absence of species 0, the RY of species from the community is given by

$$\eta_{i_{/0}} = 1 + \sum_j a_{ij}\eta_{j_{/0}}.$$

Once introduced, the RY of species 0 is given by

$$\eta_0 = 1 + \sum_i a_{0i}\eta_{i_{/0}} + \sum_i a_{0i}\Delta\eta_i.$$

where

$$\Delta\eta_i = \sum_j V_{ij}^{/0} a_{j0}\eta_0.$$

Then we have that

$$\eta_0(1 - \sum_{ij} a_{0i}V_{ij}^{/0} a_{j0}) = 1 + \sum_i a_{0i}\eta_{i_{/0}}$$

We recognize that the RHS of the equation above gives us the invasion fitness of species 0, that we note $f_0$. How species relative yield varies with its fitness is given $V_{00}$, that is

$$\eta_0 = V_{00}f_0.$$

Therefore, by identification

$$V_{00}^{-1} = 1 - \sum_{ij} a_{0i}V_{ij}^{/0} a_{j0}.$$

We simplify this equation by assuming that all species perceives an equivalent interaction from species 0 given by $\mu_0$.

$$\begin{aligned}
V_{00}^{-1} &= 1 - \mu_0 \sum_i a_{0i} \sum_j V_{ij}^{/0} \\
&= 1 - \mu_0 \sum_i a_{0i}\eta_{i_{/0}} \\
&= 1 - \mu_0(\frac{\eta_0}{V_{00}} - 1).
\end{aligned}$$

Next, we can isolate $V_{00}$ which yields

$$V_{00} = \frac{1 + \mu_0\eta_0}{1 + \mu_0}. \tag{S2}$$

We test this prediction in Fig. S4.

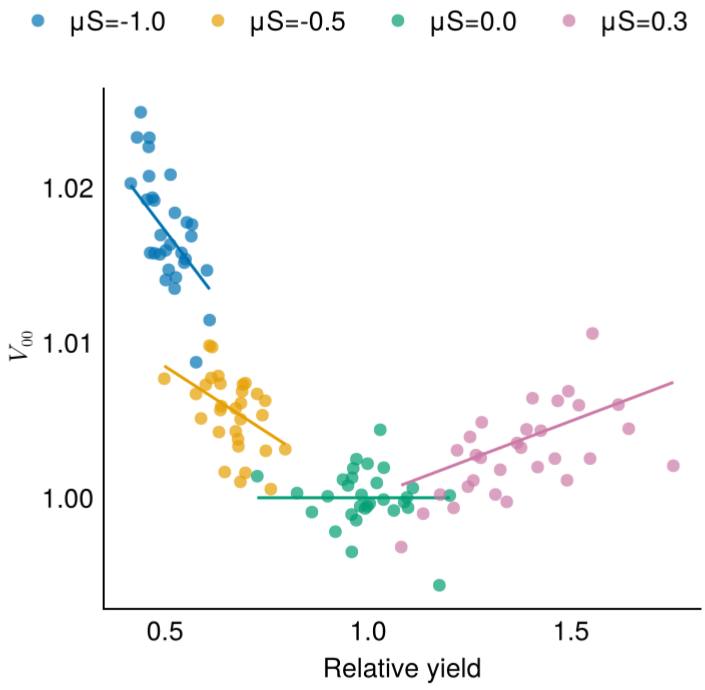


Figure S4: Species feedback against their relative yield. We simulate communities with different interactions. Each dot corresponds to the observed species feedback ($V_{00}$), and each line correspond to the analytical prediction for a $\mu_0$ assumed to be the same for all species (given by the mean of the gaussian in which interactions are drawn) (Eq. S2). Community parameters: $S = 50$, $\sqrt{S}\sigma = 0.1$, $\text{std}(K_i) = 0.3$, $r_i = 1$.

### S5 Extending predictions to smaller communities with strong interactions

Until now, we have tested our predictions in large, disordered communities characterized by relatively weak interspecific interactions. This setup is relevant to describes communities rich communities dominated by self-regulation such as in (Saavedra et al. 2017). However, some natural communities often stronger biotic interactions, and the empirical dataset we analyze comprises communities with at most six species. This raises an important question: Can our framework be applied to smaller communities with stronger interactions?

Crucially, our framework does not rely on the assumption of large species pools or vanishingly weak interspecific interactions. Instead, it is grounded in two key assumptions: (1) species reach a steady state; (2) the strength of biotic interactions is weakly correlated with the intensity of external perturbations. Regarding (1), we explain section S8 how results of previous studies could extend our work to nonequilibrium dynamics.

Specifically, the second assumption translates as assuming that covariances between biotic and disturbance impacts are close to zero. In the case of pulse disturbances, it implies $\mathrm{cov}(a_{ij}B_j^{\mathrm{eq}}, z_j) \approx 0$, and for press disturbances, $\mathrm{cov}(V_{ij}K_j, \kappa_i) \approx 0$. As interactions grow stronger, these covariances could potentially increase—but only if interactions (or disturbances) exhibit structured patterns. In the absence of such structure, we expect these correlations to remain low.

To test this, we applied our framework to communities with stronger interactions. We sampled interactions from a distribution with mean $S\mu = -2$ and standard deviation $\sqrt{S}\sigma = 2$, starting from a pool of 30 species. This places in the phase where gLV model exhibits multiple attractors (Bunin 2017). Given the strength of interactions, few species are expected to coexist. We assembled communities by simulating their dynamics until equilibrium and removed extinct species. Only communities with at least 10 surviving species were retained, ensuring sufficient data for validation. We evaluated our predictions on three such randomly assembled communities and visualized their interaction matrices, confirming that these are not dominated by self-regulation (i.e., the diagonal), see Fig. S5.

We find that while the prediction is somewhat less accurate in these smaller, strongly interacting communities, it still provides a good approximation of species sensitivity. The remaining variation can be explained by non-zero covariance between interactions and disturbance intensity—an effect that becomes more likely as interactions strengthen—and by species-specific feedbacks that deviate from unity ($V_{ii} \neq 1$). Accounting for these factors of variability is let for future studies.

Another potential source of bias in our predictions arises from changes in the steady state, particularly species extinctions. Extinctions introduce nonlinear dynamics that fall outside the assumptions of our linear framework. However, we believe that our approach could still offer insights in such cases. Specifically, under press disturbances, the species most likely to go extinct are those with the highest predicted sensitivity—indicating that our framework may help anticipate extinction risk. This represents a promising avenue for future research.

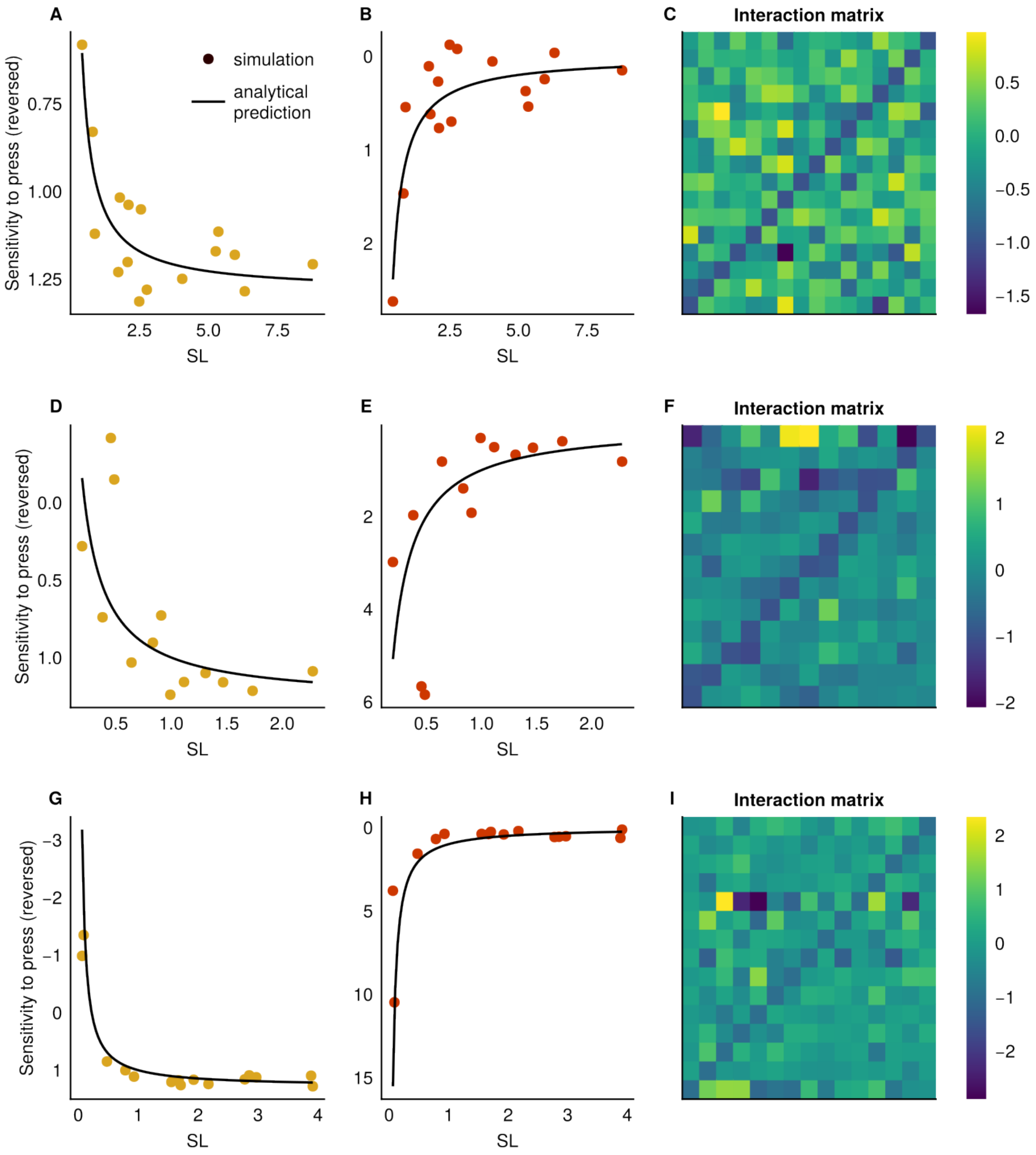


Figure S5: Same set up as Fig. 2 of the main text, with $\sqrt{S}\sigma = 2$ and $S\mu = -2$.

### S6 From competition to facilitation: sensitivity to mean interaction strengths

We tested whether our predictions hold across different types of species interactions, ranging from purely competitive to increasingly facilitative (Fig. S6). While we explore a wide spectrum of interaction strengths, there are natural limits: communities dominated by strong positive interactions are difficult to assemble because they often lead to unbounded, unrealistic population growth. Despite this, our results remain robust across the full range of feasible interaction strengths. This suggests that our framework is not limited to strictly competitive communities—it also applies to systems with facilitative interactions, such as those found in the empirical dataset we analyse in the main text (Pennekamp et al. 2018).

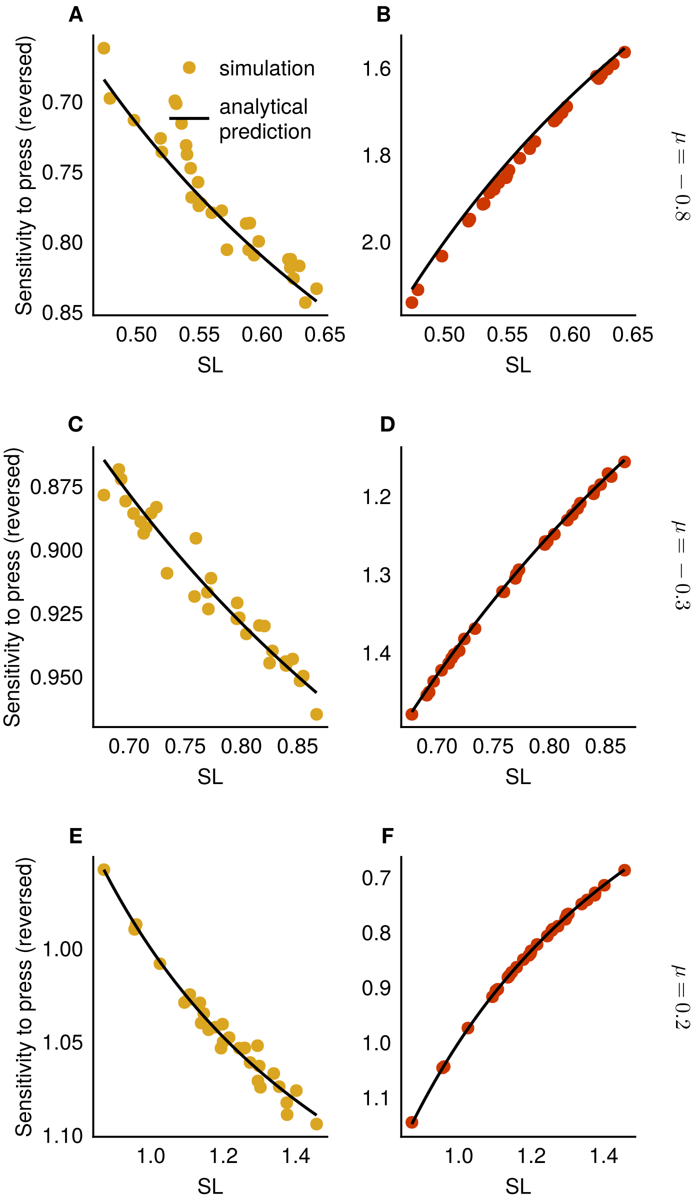


Figure S6: Effect of mean interaction strength on species' response to press disturbances. We test whether our predictions hold across communities with different average interaction strengths: strongly competitive (mean interaction $S\mu = -0.8$, panels A–B), moderately competitive ($S\mu = -0.3$, panels C–D), and weakly facilitative ($S\mu = 0.2$, panels E–F). Each community contains $S = 30$ species. We simulate 1,000 press disturbances, where the change in intrinsic growth rate is drawn from a log-normal distribution: $\delta u_i \sim \text{LogNormal}(-1, 0.5)u_i$. Community parameters: $S = 30$, $\sqrt{S}\sigma = 0.1$, $\text{std}(K_i) = 0.3$, $r_i = 1$.

### S7 Single species press: how other species respond

In the main text, we study how a species affected by targeted press (only the species of interest is affected) respond to that press. However, we could also ask how the other species, not directly affected by the press, respond. This subsection aims at answering this question.

We note species 0 the one targeted by the targeted press, and $i$ other species of the community.

The change in biomass of species not directly affected by the press write

$$\begin{aligned}\Delta B_i^{\text{eq}} &= -V_{i0}\Delta K_0 \\ &= -V_{i0}\kappa_0 K_0\end{aligned}$$

where $\kappa_0$ denotes the intensity of the press. We consider a reduction of carrying capacity, hence the minus sign in the equation above.

Then, the relative change in biomass normalized by the press intensity is given by

$$\frac{1}{\kappa_0}\frac{\Delta B_i^{\text{eq}}}{B_i^{\text{eq}}} = -V_{i0}\frac{K_0}{B_i^{\text{eq}}}$$

At this point, we cannot say much because the response of species $i$ is depends on its specific relationship with species 0. What we can do is ask a more general question: how species $i$ respond to targeted press on another species *on average*?

To answer this question, we have to consider the average of the quantity above, which will allows us to simplify it

$$\langle\frac{1}{\kappa_0}\frac{\Delta B_i^{\text{eq}}}{B_i^{\text{eq}}}\rangle_{0\neq i} = -\frac{1}{S-1}\frac{1}{B_i^{\text{eq}}}\sum_{0\neq i} V_{i0}K_0$$

Here, we can use that

$$\sum_{0\neq i} V_{i0}K_0 = B_i^{\text{eq}} - V_{ii}K_i$$

which yields

$$\langle\frac{1}{\kappa_0}\frac{\Delta B_i^{\text{eq}}}{B_i^{\text{eq}}}\rangle_{0\neq i} = \frac{1}{S-1}(\frac{V_{ii}}{\text{SL}_i} - 1) \tag{S3}$$

We recover a similar pattern than for the community wide-press: species stability increases with their self-regulation. That is, because in this setting species response to the press is only determined by the interaction release. Facilitated species (SL $>$ 1) will on average be subjected to a decrease in biomass (assuming $V_{ii}$ close to one). Conversely, outcompeted species (SL $<$ 1) will see their biomass increase.

We validate this prediction in Fig. S7.

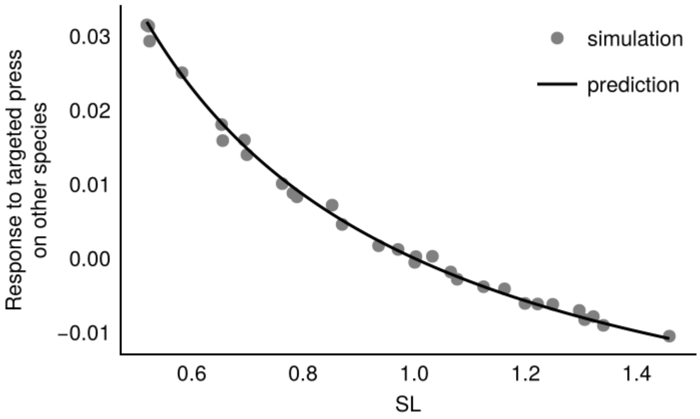


Figure S7: Response of *other* species to a targeted press. We apply to targeted press on a single species and measure the response of others. We repeat this for all species of the community, and measure the average response. Prediction based on Eq. S3. Community parameters: $S = 30$, $S\mu = 0$, $\sqrt{S}\sigma = 0.3$, $\mathrm{std}(K_i) = 0.3$, $r_i = 1$, $\kappa = 0.1$.

### S8 Link with Medeiros et al. 2023

Medeiros et al. 2023 investigate how biotic interactions influence species sensitivity in systems with non-equilibrium dynamics. Although their definition of sensitivity differs from ours due to this focus, potential links between the two frameworks exist, and we explore them here.

In their approach, species sensitivity is defined as the distance between perturbed and unperturbed trajectories over time. A species is considered highly sensitive if a small perturbation is amplified over time, and less sensitive if it is dampened. In contrast, we focus on systems at ecological equilibrium, where we define species sensitivity as the degree to which a change in environmental conditions leads to an amplified or dampened change in biomass at equilibrium. This definition does not involve temporal dynamics directly.

However, our results reveal that responses to press (environmental change) and pulse (biomass removal) disturbances are closely linked via self-regulation loss (SL). Since pulse responses inherently include a temporal component, this invites comparison with the non-equilibrium perspective of Medeiros et al. 2023.

In particular, Medeiros et al. 2023 show that species sensitivity can be approximated by their contribution to the dominant eigenvector of the community matrix. The dominant eigenvalue governs the long-term recovery rate of the system, and a species' corresponding eigenvector coefficient indicates its contribution to that recovery. A coefficient near zero implies that the species rapidly returns to equilibrium and does not limit community recovery, while a coefficient near one (assuming the eigenvector is normalized) indicates that the species is strongly aligned with the slowest mode of recovery, and thus limits the system's return to equilibrium.

Building on this idea, they propose that the magnitude of a species' alignment with the dominant eigenvector, $|v_{1i}|$, provides a useful proxy for its sensitivity.

To explore the connection with our framework, we examined the relationship between species alignment $|v_{1i}|$ and their SL in 10 randomly assembled communities (Fig. S8). We found a clear, negative relationship: species most strongly aligned with the dominant eigenvector tend to have low SL. This finding aligns with (Arnoldi et al. 2018), who showed that species with low biomass relative to their carrying capacity—equivalent to SL $\ll 1$ in our framework—are typically the slowest to recover from disturbances.

Taken together, these results suggest that our work and that of Medeiros et al. 2023 are complementary. While our study provides a mechanistic and equilibrium-based understanding of how biotic interactions shape species sensitivity, their framework shows that similar sensitivity rankings can persist beyond equilibrium, opening a promising direction for bridging stability concepts across dynamic regimes.

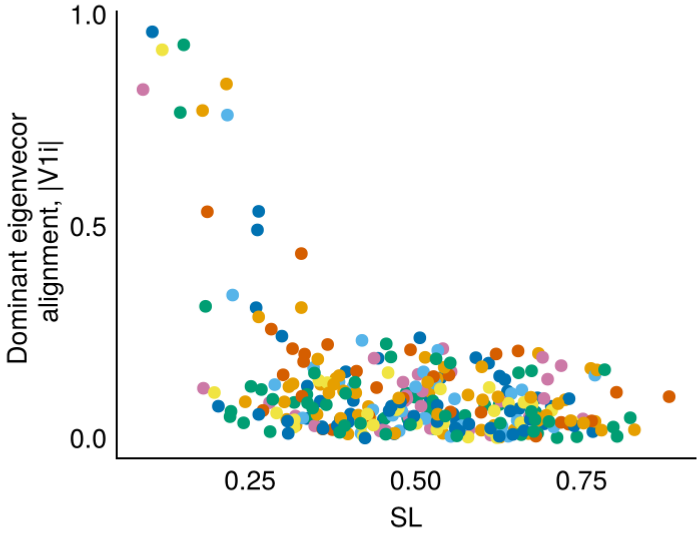


Figure S8: Species' alignment with eigenvector (sensu (Medeiros et al. 2023) against their SL for 10 random communities. Color correspond to different communities. Community parameters: $S = 30$, $S\mu = -1$, $\sqrt{S}\sigma = 0.3$, $\mathrm{std}(K_i) = 0.3$, $r_i = 1$.

### S9 Heterogeneity in species growth rates

We investigate how heterogeneity in species growth rates ($r_i$) affects our results. Using the same simulation setting as in Fig. 3 (main text), we now draw growth rates from a uniform distribution: $r_i \sim \text{Uniform}(0.1, 2)$. Results are shown in Fig. S9. We find that our predictions remain accurate in the short term but degrade over time. This suggests that heterogeneity in growth rates introduces variability through the interaction of species operating on different timescales—a divergence that accumulates with time. Understanding how biotic interactions and growth-rate heterogeneity jointly shape species stability is an important direction for future research.

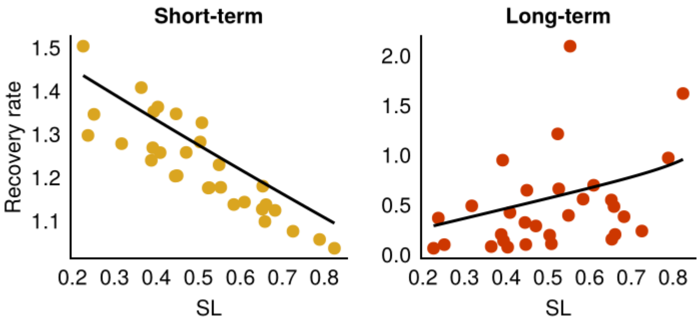


Figure S9: Species recovery rates under heterogeneous growth rates. Short-term responses are measured at $t = 0.1$, and long-term responses at $t = 10$. To account for variation in intrinsic growth rates, recovery rates are normalized by each species' own $r_i$ (i.e., observed recovery rate divided by $r_i$).

### References

Arnoldi, J. -F. et al. (Jan. 2018). “How Ecosystems Recover from Pulse Perturbations: A Theory of Short- to Long-Term Responses”. In: *Journal of Theoretical Biology* 436, pp. 79–92. ISSN: 0022-5193. DOI: 10.1016/j.jtbi.2017.10.003.

Bunin, Guy (Apr. 2017). “Ecological Communities with Lotka-Volterra Dynamics”. In: *Physical Review E* 95.4, p. 042414. DOI: 10.1103/PhysRevE.95.042414.

Ives, A. R., K. Gross, and J. L. Klug (Oct. 1999). “Stability and Variability in Competitive Communities”. In: *Science* 286.5439, pp. 542–544. DOI: 10.1126/science.286.5439.542.

Loreau, Michel and Claire de Mazancourt (2013). “Biodiversity and Ecosystem Stability: A Synthesis of Underlying Mechanisms”. In: *Ecology Letters* 16.s1, pp. 106–115. ISSN: 1461-0248. DOI: 10.1111/ele.12073.

Loreau, Michel and Andy Hector (July 2001). “Partitioning Selection and Complementarity in Biodiversity Experiments”. In: *Nature* 412.6842, pp. 72–76. ISSN: 1476-4687. DOI: 10.1038/35083573.

Medeiros, Lucas P. et al. (2023). “Ranking Species Based on Sensitivity to Perturbations under Non-Equilibrium Community Dynamics”. In: *Ecology Letters* 26.1, pp. 170–183. ISSN: 1461-0248. DOI: 10.1111/ele.14131.

Mentges, Andrea et al. (2024). “Accounting for Effects of Growth Rate When Measuring Ecological Stability in Response to Pulse Perturbations”. In: *Ecology and Evolution* 14.10, e11637. ISSN: 2045-7758. DOI: 10.1002/ece3.11637.

Pennekamp, Frank et al. (Nov. 2018). “Biodiversity Increases and Decreases Ecosystem Stability”. In: *Nature* 563.7729, pp. 109–112. ISSN: 1476-4687. DOI: 10.1038/s41586-018-0627-8.

Saavedra, Serguei et al. (2017). “A Structural Approach for Understanding Multispecies Coexistence”. In: *Ecological Monographs* 87.3, pp. 470–486. ISSN: 1557-7015. DOI: 10.1002/ecm.1263.

Sibly, Richard M. et al. (July 2005). “On the Regulation of Populations of Mammals, Birds, Fish, and Insects”. In: *Science* 309.5734, pp. 607–610. DOI: 10.1126/science.1110760.
